# Site-specific amino-acid preferences are mostly conserved in two closely related protein homologs

**DOI:** 10.1101/018457

**Authors:** Michael B. Doud, Orr Ashenberg, Jesse D. Bloom

## Abstract

Evolution drives changes in a protein’s sequence over time. The extent to which these changes in sequence lead to shifts in the underlying preference for each amino acid at each site is an important question with implications for comparative sequence-analysis methods such as molecular phylogenetics. To quantify the extent that site-specific amino-acid preferences shift during evolution, we performed deep mutational scanning on two homologs of human influenza nucleoprotein with 94% amino-acid identity. We found that only a modest fraction of sites exhibited shifts in amino-acid preferences that exceeded the noise in our experiments. Furthermore, even among sites that did exhibit detectable shifts, the magnitude tended to be small relative to differences between non-homologous proteins. Given the limited change in amino-acid preferences between these close homologs, we tested whether our measurements could inform site-specific substitution models that describe the evolution of nucleoproteins from more diverse influenza viruses. We found that site-specific evolutionary models informed by our experiments greatly outperformed non-site-specific alternatives in fitting phylogenies of nucleoproteins from human, swine, equine, and avian influenza. Combining the experimental data from both homologs improved phylogenetic fit, partly because measurements in multiple genetic contexts better captured the evolutionary average of the amino-acid preferences for sites with shifting preferences. Our results show that site-specific amino-acid preferences are sufficiently conserved that measuring mutational effects in one protein provides information that can improve quantitative evolutionary modeling of nearby homologs.

## Introduction

Since the first comparative analyses of homologous proteins by Zuckerkandl and Pauling (1965) fifty years ago, it has been obvious that different sites in proteins evolve under different constraints, with some sites substituting to a wide range of amino acids, while others are constrained to one or a few identities. Zuckerkandl and Pauling (1965) proposed, and decades of subsequent work have confirmed (De-Pristo et al., 2005; Harms and Thornton, 2013), that these constraints arise from the highly cooperative interactions among sites that shape important protein properties such as stability, folding kinetics, and biochemical function.

The complexity and among-sites cooperativity of these evolutionary constraints mean that a mutation at a single site can in principle shift the amino-acid preferences of any other site – and numerous experiments have demonstrated examples of such epistasis among sites (Weinreich et al., 2006; Ortlund et al., 2007; da Silva et al., 2010; Lunzer et al., 2010; Gong et al., 2013; Natarajan et al., 2013; Podgornaia and Laub, 2015). However, experiments have also shown that despite such epistasis, the amino-acid preferences of many sites are similar across homologs (Risso et al., 2015; Ashenberg et al., 2013; Serrano et al., 1993). For instance, protein structures themselves are highly conserved during evolution (Chothia and Lesk, 1986; Sander and Schneider, 1991), and sites in specific structural contexts often have strong propensities for certain amino acids (Chou and Fasman, 1974; Richardson and Richardson, 1988; Lim and Sauer, 1991). Furthermore, many of the most successful methods for identifying distant homologs (e.g. PSI-BLAST) utilize site-specific scoring models (Henikoff and Henikoff, 1997; Altschul et al., 1997), implying that amino-acid preferences are at least somewhat conserved even among homologs with low sequence identity.

A half-century of work has therefore made it abundantly clear that site-specific amino-acid preferences can in principle shift arbitrarily during evolution, but nonetheless in practice remain somewhat conserved among homologs. The important remaining question is the *extent* to which site-specific aminoacid preferences are conserved versus shifted. This question is especially important for the development of quantitative evolutionary models for tasks such as phylogenetic inference. Initially, phylogenetic models unrealistically assumed that sites within proteins evolved both independently and under identical constraints. But more recent models have relaxed the second assumption that sites evolve identically. At first, this relaxation only allowed sites to evolve at different rates (Yang, 1994). But newer models also accommodate variation in the amino-acid preferences among sites, either by treating these preferences as parameters of the substitution model (Lartillot and Philippe, 2004; Le et al., 2008; Wang et al., 2008; Rodrigue et al., 2010) or by leveraging their direct measurement by high-throughput experiments (Bloom, 2014a,b). Because these models retain the assumption of independence among sites, they will outperform traditional non-site-specific models only if site-specific amino-acid preferences are substantially conserved among homologs.

Here we perform the first experimental quantification of the conservation of the amino-acid preferences at all sites in two homologous proteins. We do this by using deep mutational scanning (Fowler et al., 2010; Fowler and Fields, 2014) to comprehensively measure the effects of all mutations to two homologs of influenza nucleoprotein (NP) with 94% sequence identity. We find that the amino-acid preferences are substantially conserved at most sites in the homologs, but some sites have significant shifts in preferences. We then test whether the experimentally measured site-specific amino-acid preferences can inform site-specific phylogenetic substitution models that describe the evolution of more diverged NP homologs. We find that the experimentally informed site-specific substitution models exhibit improved fit to NP phylogenies containing diverged sequences from human, swine, equine, and avian influenza lineages. Overall, our work shows that site-specific amino-acid preferences are sufficiently conserved that measurements on one homolog can be used to improve the quantitative evolutionary modeling of closely related homologs.

## Results

### Comparison of amino-acid preferences between two homologs

#### Deep mutational scanning of two influenza NP homologs

Our studies focused on NP from influenza A virus. NP performs several conserved functions that are essential for the viral life cycle, including encapsidation of viral RNA into ribonucleoprotein complexes for transcription, viral-genome replication, and viral-genome trafficking (Eisfeld et al., 2015). NP’s structure is highly conserved in all characterized influenza strains (Ye et al., 2006; Das et al., 2010). Our studies compared the site-specific amino-acid preferences of NP homologs from two human influenza strains, PR/1934 (H1N1) and Aichi/1968 (H3N2) (Figure 1). These NPs have diverged by over 30 years of evolution, and differ at 30 of their 498 residues (94% protein sequence identity).

**Figure 1:**
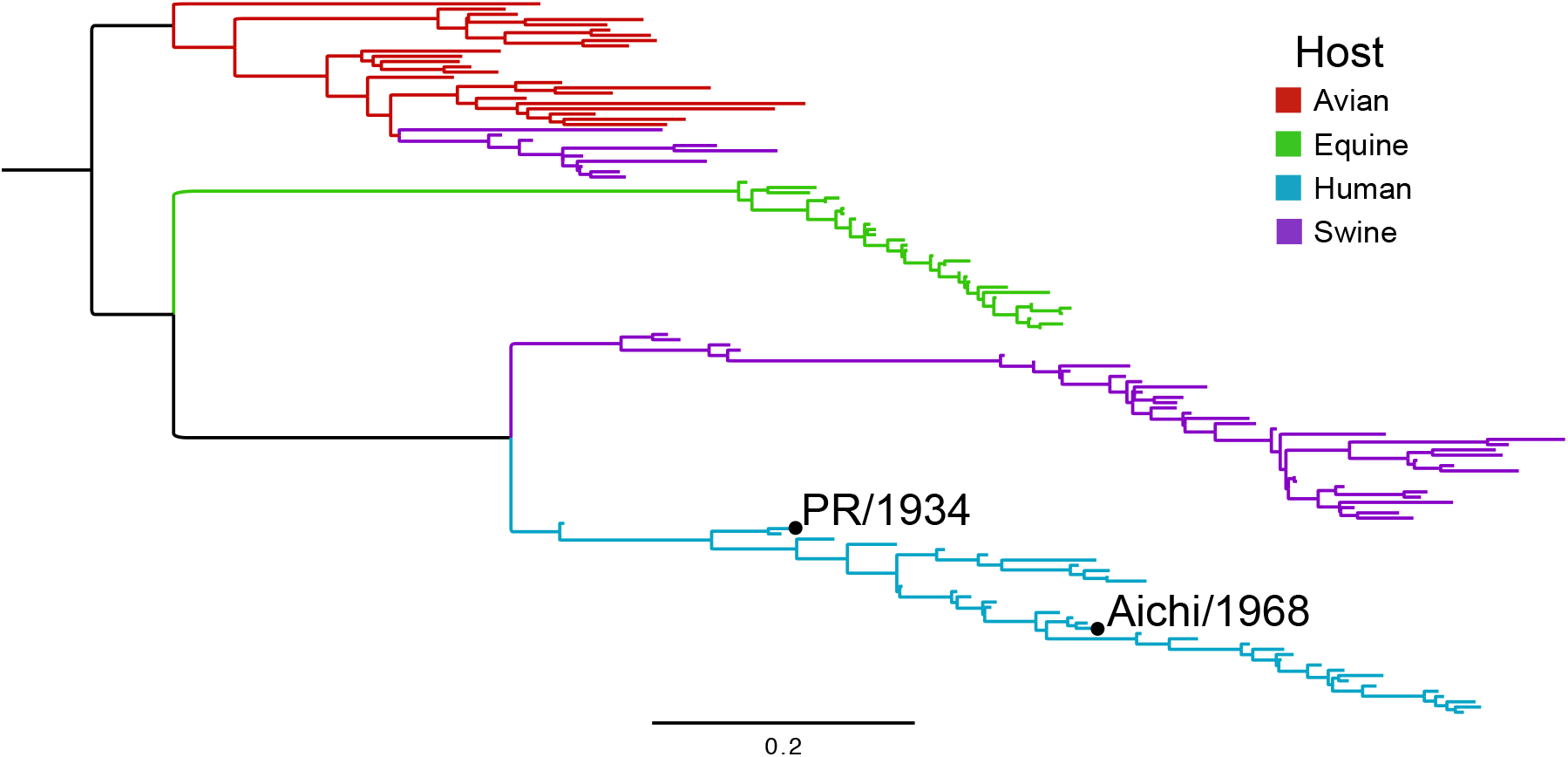
Phylogenetic tree of influenza NPs. The two homologs used in this work are labeled on the human influenza lineage. A diverse set of sequences was collected by sampling across years and hosts, and a maximum-likelihood tree was inferred using `CodonPhyML` (Gil et al., 2013) with the codon substitution model of Goldman and Yang (1994). The tree was rooted using the avian clade as an outgroup. The scale bar is in units of codon substitutions per site.

We used our previously described approach for deep mutational scanning of influenza genes (Bloom, 2014a; Thyagarajan and Bloom, 2014) to measure the site-specific amino acid preferences of the PR/1934 and Aichi/1968 NPs. Briefly, this approach involved using a PCR-based technique to create mutant libraries of plasmids encoding NP genes with random codon mutations, using reverse genetics to incorporate these mutant genes into influenza viruses, and then passaging these viruses at low multiplicity of infection to select for viruses carrying functional NP variants. Deep sequencing was used to count the occurrences of each mutation before and after selection, and the amino-acid preferences for each site were inferred from these counts using `dms_tools` (Bloom, 2015) (Supplementary file 1, Supplementary file 2, Supplementary figure 1, Supplementary figure 2). Our mutagenesis randomized 497 of the 498 codons in NP (the N-terminal methionine was not mutagenized), and so our libraries sampled all 497 *×* 19 = 9, 443 amino-acid mutations at these sites. Our mutagenesis introduced an average of about two codon mutations per gene, with the number of mutations per gene following a roughly Poisson distribution (Supplementary figure 3), and so the effect of each mutation was assayed both alone and in the background of variants that contained one or more additional mutations.

Because deep mutational scanning is subject to substantial experimental noise, we performed several full biological replicates for each NP homolog, beginning with independent creation of the plasmid mutant library. In the current work, we performed three replicates of deep mutational scanning on the PR/1934 NP and two replicates on the Aichi/1968 NP. In a previous study (Bloom, 2014a) we performed eight replicates of deep mutational scanning on Aichi/1968 NP. We will refer to these previous replicates of the Aichi/1968 NP deep mutational scanning as the *previous study*, and the two new replicates as the *current study*. When not otherwise noted, we refer to the pooled data of all ten of these replicates simply as *Aichi/1968*.

#### Amino-acid preferences are well correlated between homologs

For each homolog we averaged the site-specific amino-acid preferences across all replicates and examined the correlations of the preferences for each of the 20 amino acids at each of the 497 sites we mutagenized (all sites can be unambiguously aligned between homologs). The mean preferences for the two NP homologs have a Pearson’s correlation coefficient of 0.78 (Figure 2A). In comparison, the correlation between the preferences measured in the *previous study* and *current study* on the Aichi/1968 homolog is 0.83 (Figure 2B). Therefore, the amino-acid preferences correlate nearly as well between the two homologs as they do between different experiments on the same homolog. As expected, there is no correlation between the preferences of the PR/1934 NP and a non-homologous protein (hemagglutinin, HA) for which we have previously measured the site-specific amino-acid preferences using the same approach as in this work (Thyagarajan and Bloom, 2014) (Figure 2C).

**Figure 2:**
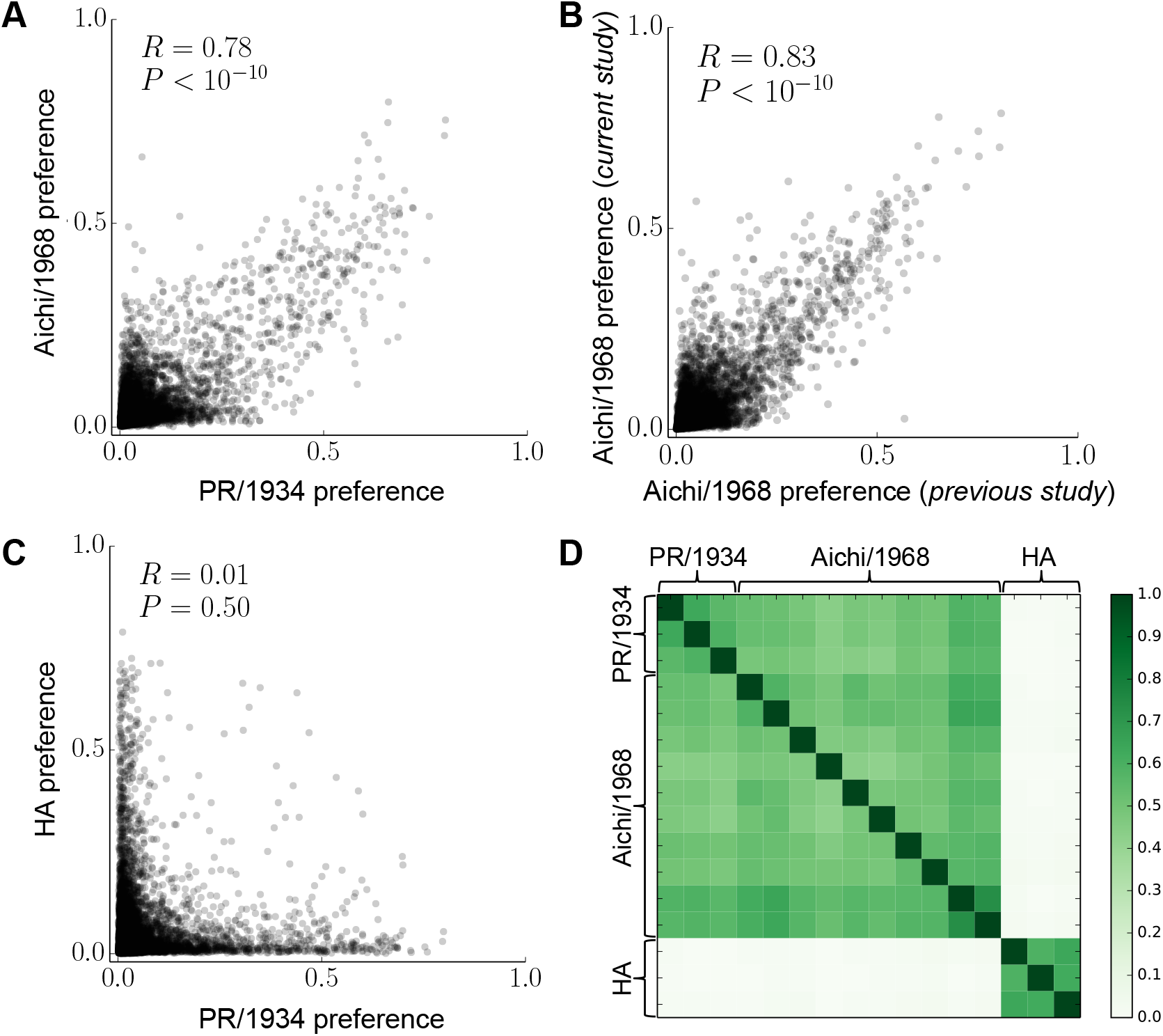
Site-specific amino-acid preferences correlate nearly as well between NP homologs as between replicate measurements on the same homolog. (A), (B) The correlation between the mean of the preferences taken over all replicates on each NP homolog is nearly as large as that between the preferences measured in the *current study* and *previous study* on the Aichi/1968 NP. (C) However, there is no correlation between the preferences measured for NP and the non-homologous protein HA. Each data point in (A)-(C) is the preference for one of the twenty amino acids at one of the 497 sites in NP. *R* is the Pearson correlation coefficient. (D) The Pearson correlations between the preferences measured in all pairs of individual replicates. Comparisons between NP and HA were made based on position in primary sequence for sites 2 through 498.

We also asked if the site-specific amino-acid preferences from each replicate showed the same pattern of correlation between homologs that we observed when comparing mean preferences. We again found that correlation coefficients are just as high between NP homologs as they are between replicate measurements on the same homolog, and that there is no correlation between the preferences for NP and the non-homologous protein HA (Figure 2D). Overall, these results indicate that at the vast majority of sites, any differences in the amino-acid preferences between NP homologs are smaller than the noise in our experimental measurements, and vastly smaller than the differences between non-homologous proteins.

#### Shifts in amino-acid preferences are small for most sites

The previous section shows that any widespread shifts in site-specific amino-acid preferences are smaller than the noise in our experiments. However, it remains possible that a subset of sites show substantial shifts in their amino-acid preferences that are masked by examining all sites together. We therefore performed an analysis to identify specific sites with shifted amino-acid preferences between homologs.

This analysis needed to account for the fact that experimental noise induced variation in the prefer-ences measured in each replicate. Figure 3 shows replicate measurements for both homologs at several sites in NP. At many sites, such as site 298, all replicate measurements yielded highly reproducible amino-acid preferences both between and within homologs. At many other sites, such as site 3, replicate measurements were quite variable both between and within homologs, probably due to fairly weak selection at that site. Some sites, like site 254, exhibited reproducible measurements within each homolog, and the most preferred amino acid was the same in both homologs, but the tolerance for mutations to other amino acids was distinct in each homolog. Finally, at a few sites, most prominently site 470, replicate measurements were highly reproducible within each homolog but clearly differed in which amino acid was most preferred between homologs. We therefore developed a quantitative measure of the shift in preferences between homologs that accounts for this site-specific experimental noise.

**Figure 3:**
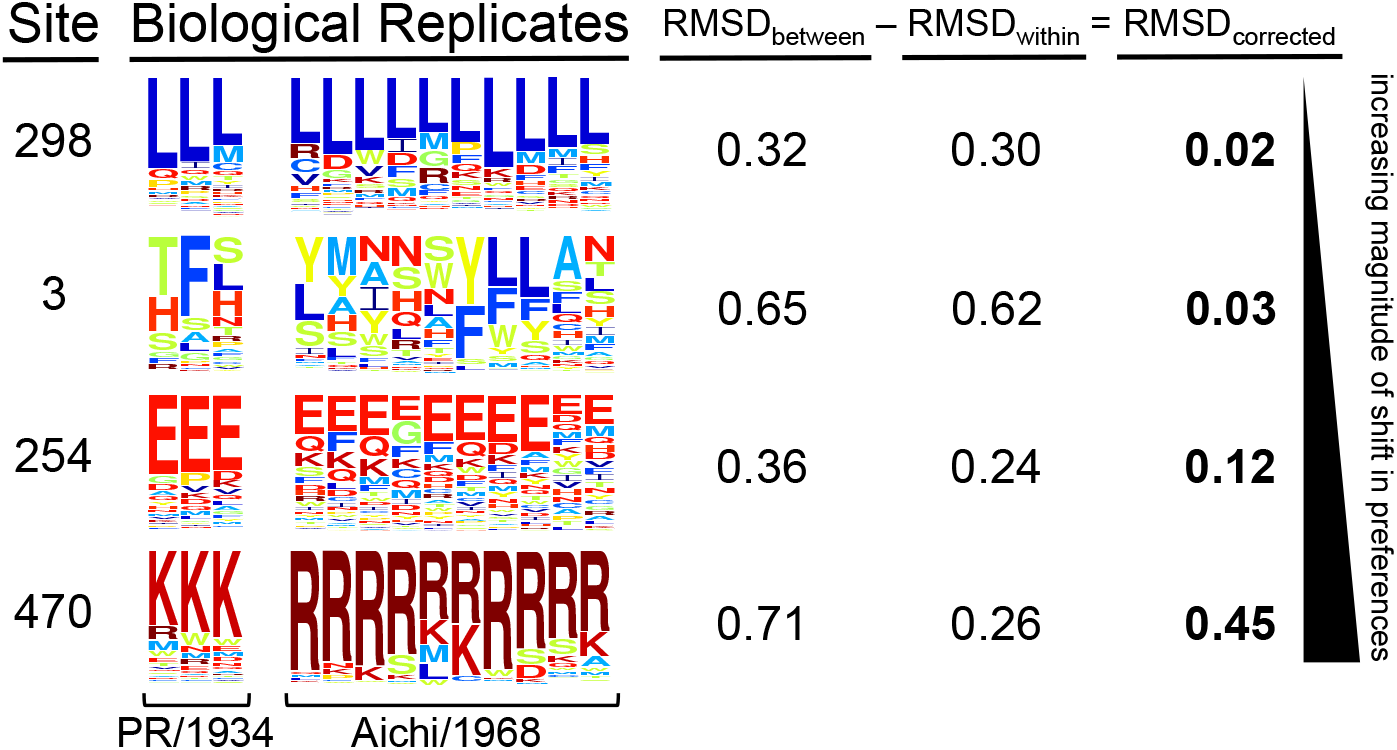
Replicate measurements quantify the shift in amino-acid preferences between homologs after correcting for experimental noise. The amino-acid preferences measured in multiple replicates of deep mutational scanning of both homologs are shown for selected sites ordered by the magnitude of preference change observed after correcting for site-specific noise. *RMSD_between_* (the average difference between the two homologs) and *RMSD_within_* (the average variation within replicates of each homolog) are shown to the right. *RMSD_corrected_* is calculated by subtracting *RMSD_within_* from *RMSD_between_*.

We used the Jensen-Shannon distance metric (the square root of the Jensen-Shannon divergence) to quantify the distance between the 20-dimensional vectors of amino-acid preferences for each pair of replicate measurements at each site. This distance ranges from zero (identical amino-acid preferences) to one (completely different amino-acid preferences). To quantify experimental noise at a site, we calculated the root-mean-square of the Jensen-Shannon distance for all pairwise comparisons among replicate measurements on the same homolog, and termed this quantity *RMSD_within_*. Sites with large *RMSD_within_* have high experimental noise. We defined an analogous statistic, *RMSD_between_*, to quantify the distance in preferences between homologs by calculating the root-mean-square of the Jensen-Shannon distance for all pairwise comparisons between replicates of PR/1934 and replicates of Aichi/1968. Figure 3 shows the values of these statistics for example sites.

The fact that we had data from two independent sets of experiments on the Aichi/1968 NP (the *current study* and *previous study*) enabled us to perform a control analysis by calculating *RMSD_between_* and *RMSD_within_* for the replicates from these two experiments. As an additional control to gauge the extent of amino-acid preference differences between non-homologous proteins, we also calculated *RMSD_between_* and *RMSD_within_* for our experiments on Aichi/1968 NP and our previous study on HA (note that because NP and HA are non-homologous, they cannot be meaningfully aligned, so this control comparison simply pairs each site in NP with the corresponding residue number in HA).

The relationship between *RMSD_between_* (the observed difference between homologs) and *RMSD_within_* (the observed variation in repeated measurements on the same homolog) for all sites is shown for several different comparisons in Figure 4A-C. Sites with low *RMSD_within_* exhibit highly reproducible measurements between replicate experiments, whereas sites with higher values of *RMSD_within_* exhibit substantial experimental noise, probably due to weak selection at that site. Sites with large *RMSD_between_* exhibit amino-acid preference differences between homologs, but at each site some of this observed variation is due to the site-specific experimental noise (quantified by *RMSD_within_*) rather than a true difference between the homologs.

**Figure 4:**
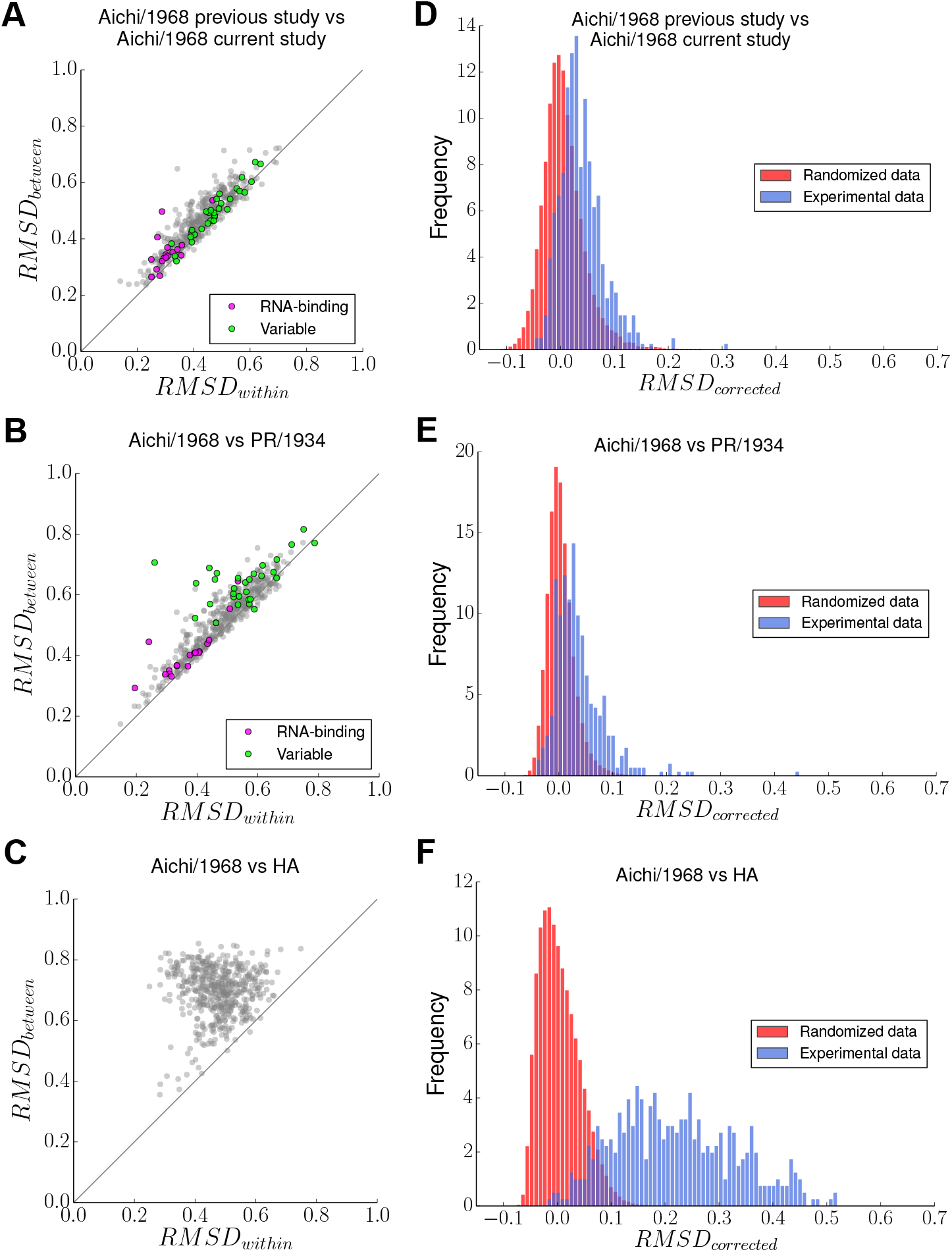
Identification of sites with shifts in amino-acid preferences. (A)-(C) Each plot shows statistics calculated for a comparison between two groups of replicate experiments. Each point represents a site in NP. *RMSD_within_* quantifies the average difference in amino-acid preferences within each of the two groups (experimental noise), and *RMSD_between_* quantifies the average difference in preferences between the two groups. Points above the *y* = *x* diagonal represent sites with preference changes between homologs greater than experimental noise. Sites in the RNA-binding groove are in purple; sites that have different wild-type identities in PR/1934 and Aichi/1968 are in green. (D)-(F) The actual distribution of *RMSD_corrected_* values is shown in blue, and the distribution of *RMSD_corrected_* from data randomized between comparison groups is shown in red. Comparisons are made between the two studies on Aichi/1968 (A, D), between Aichi/1968 and PR/1934 (B, E), and between Aichi/1968 and the non-homologous HA (C, F).

When comparing two independent experiments on the same NP (Figure 4A) or comparing experiments on two homologs of NP (Figure 4B), the relationship between *RMSD_between_* and *RMSD_within_* is approximately linear, indicating that the difference in amino-acid preferences between homologs at a given site is usually comparable to the experimental noise. Deviations from this linear relationship are more frequent in the comparison between PR/1934 and Aichi/1968 (Figure 4B) than in the comparison between the two studies of Aichi/1968 (Figure 4A). These deviations mostly arise from sites that have larger *RMSD_between_* than *RMSD_within_*, indicating that these sites have shifts in their amino-acid preferences between homologs that exceed the experimental noise. These results comparing NP homologs are in stark contrast with the *RMSD_between_* and *RMSD_within_* calculated when comparing NP to the non-homologous HA (Figure 4C), where the difference between proteins is almost always substantially greater than the experimental noise.

To quantify the extent of amino-acid preference shifts between the two homologs in a way that corrects for the experimental noise, we defined another statistic, *RMSD_corrected_*, by subtracting *RMSD_within_* from *RMSD_between_* (Figure 3, Supplementary file 5). Sites with shifts in amino-acid preferences greater than the experimental noise have *RMSD_corrected_ >* 0. However, we also expect many sites to have positive *RMSD_corrected_* values due to statistical noise. To determine the distribution of *RMSD_corrected_* values expected due to such statistical noise alone under the null hypothesis that the amino-acid preferences are the same in both groups being compared, we generated null distributions of *RMSD_corrected_* using exact randomization testing by shuffling which experimental replicates were assigned to which NP homolog. For every possible shuffling of replicates, we computed *RMSD_corrected_* at every site and combined the results across all shufflings.

The distribution of *RMSD_corrected_* obtained experimentally mostly overlaps the randomized distribution of *RMSD_corrected_* when comparing the two independent Aichi/1968 experiments (Figure 4D). This overlap is consistent with the hypothesis that the true amino-acid preferences are the same in both experiments on the Aichi/1968 NP. In contrast, when comparing PR/1934 to Aichi/1968, some *RMSD_corrected_* values are shifted in the positive direction substantially beyond the null distribution (Figure 4E), indicating larger differences in preferences at some sites than can be explained by experimental noise alone. This shift in preferences is particularly notable for site 470, which has a *RMSD_corrected_* of 0.45 as illustrated in Figure 3. However, most sites still fall within the null distribution when comparing the two NP homologs. In contrast, if NP is compared to the non-homologous HA, the vast majority of sites exhibit differences in preferences that vastly exceed the values expected under the null distribution (Figure 4F).

As an alternative approach to generating null distributions of *RMSD_corrected_*, we performed simulations of observed amino-acid preferences in each replicate under a model where there are no differences in the underlying preferences between the two homologs, but varying levels of noise for each experiment. We simulated amino-acid preferences at each site by drawing from a Dirichlet distribution, which is well-suited for this purpose because its support is a normalized vector of values, in this case corre-sponding to the vector of amino-acid preferences at a site. Our null hypothesis is that the amino-acid preferences are the same for both homologs, so we performed simulations assuming that the *true* vector of amino-acid preferences at a site is equal to the average of our experimental measurements for both homologs. We simulated the amino-acid preferences for each replicate by drawing from a Dirichlet distribution centered on this vector of assumed true preferences. The extent to which any given sample drawn from this Dirichlet distribution differs from the true vector can be tuned with a single scaling parameter (the concentration parameter). We identified a value for the concentration parameter for each experiment (Aichi/1968 *current study*, Aichi/1968 *previous study*, and PR/1934) that resulted in correlation coefficients between replicates that matched those in the actual experiment. We performed 1000 replicate simulations and combined the calculated *RMSD_corrected_* values from all simulations to build the null distribution. The distributions of *RMSD_corrected_* obtained by simulation are in Supplementary figure 5 and are similar to those obtained using exact randomization testing.

#### Sites with clear shifts in amino-acid preferences

Using either of the two null distributions, we were able to identify specific sites with *RMSD_corrected_* values significantly larger than expected due to experimental noise alone (Supplementary file 5). These are sites for which we can reject the null hypothesis that there is no shift in amino-acid preferences. To control for multiple hypothesis testing, we set a false discovery rate (proportion of rejected null hypotheses expected to be falsely rejected) of 5%.

Using exact randomization as a null distribution, we could reject the null hypothesis of no shift in amino-acid preference for 14 of the 497 sites. The simulated-data null distribution appeared to afford greater statistical power, and allowed us to reject the null hypothesis of no shift in preference for 76 sites (the 14 identified by the exact randomization plus an additional 62). Many of these additional sites, however, exhibit shifts that are small in magnitude; for instance, 30 of the additional 62 sites show a pattern similar to that of site 254 (Figure 3), where the most preferred amino acid is unchanged, but the tolerance for mutations to other residues is somewhat larger in one homolog than the other.

Figure 4 provides a more visual way to gauge the magnitude of the shifts in amino-acid preferences. If the preferences are completely conserved among homologs, the actual distribution in Figure 4E should look roughly like that in Figure 4D. In contrast, if the preferences have completely shifted between homologs, the actual distribution should look more like that in Figure 4F. As is clear from visual inspection, only a handful of sites have amino-acid preferences that have shifted between the PR/1934 and Aichi/1968 homologs to be as different as is typical for pairs of sites from non-homologous proteins. The rest of the sites either exhibit a more modest shift in preference (this is the case for 14 or 76 sites depending on which null distribution is used) or no detectable shift in preference.

An important question is whether there are common characteristics of sites with shifted preferences. One reasonable hypothesis is that sites with wild-type amino-acid identities that differ between the homologs are more likely to have experienced shifts in their amino-acid preferences. Among the 14 sites identified as shifted by both null distributions, 5 have different wild-type amino-acid identities in PR/1934 and Aichi/1968 (Figure 5A). Therefore, of sites with variable amino-acid identity between the two homologs, 17% exhibit clear shifts in preference identified by both null distributions, while only 2% of conserved sites exhibit comparable shifts.

**Figure 5:**
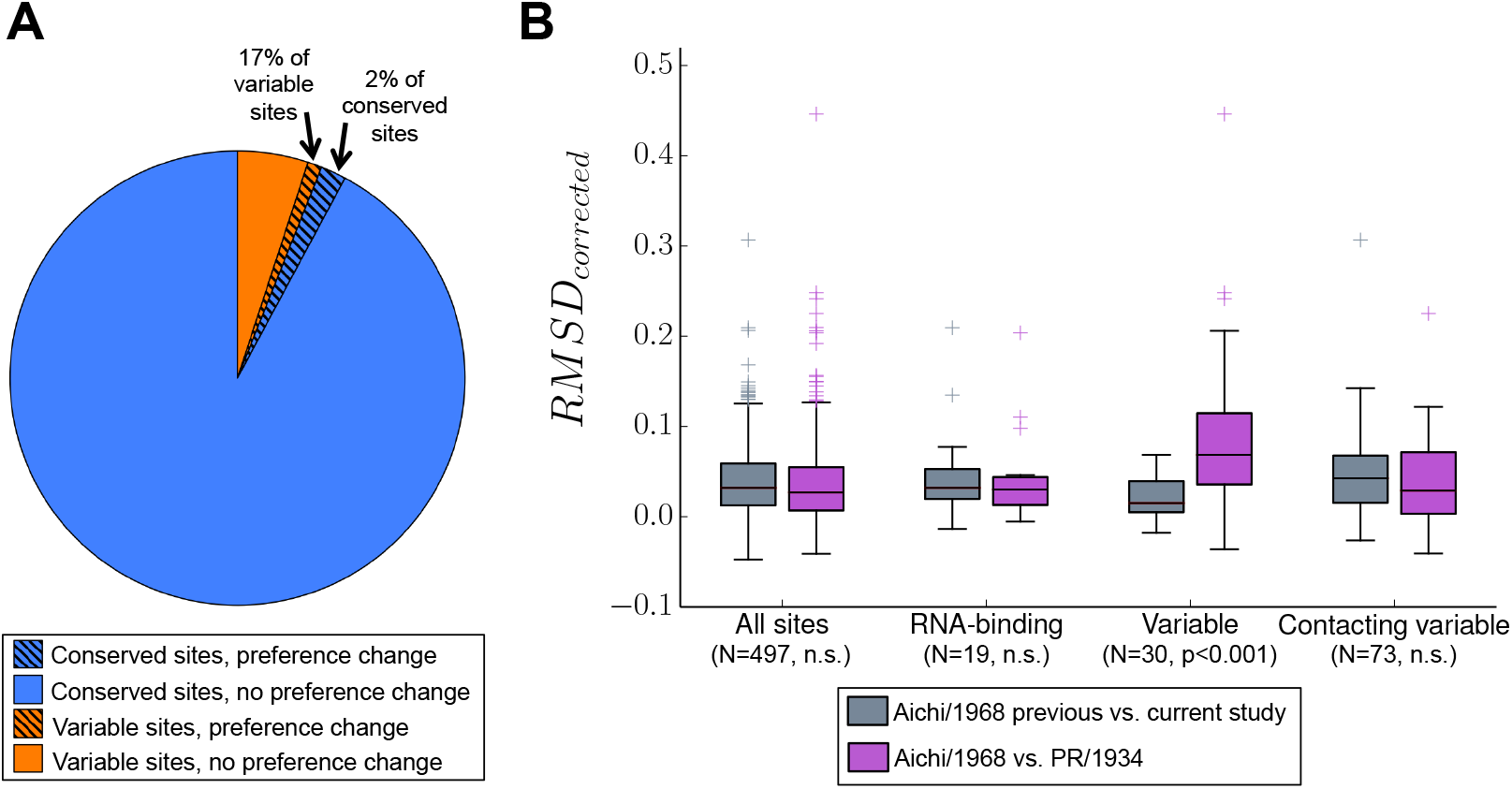
Evolutionarily variable sites are enriched for changes in amino-acid preference. (A) Sites with shifts in amino-acid preferences were identified by *RMSD_corrected_* values greater than expected under a null model assuming no difference between homologs (false discovery rate of 5% using a null model generated by exact randomization testing). *Variable* sites have different wild-type residues in the two NP homologs. (B) The distributions of *RMSD_corrected_* for various groups of sites. The median is marked by a horizontal line, boxes extend from 25th to 75th percentile, and whiskers extend to data points within 1.5 times the interquartile range. Outliers are marked with crosses. *Contacting variable* sites are conserved sites with side-chain atoms within 4.5 Ångströms of a variable side-chain atom. *RMSD_corrected_* distributions for each group of sites are shown for two comparisons: one comparing two independent experiments on Aichi/1968, and one comparing Aichi/1968 to PR/1934. P-values were determined using the Mann-Whitney U test and adjusted using the Bonferroni correction.

Having identified evolutionarily variable sites as enriched for the clearest shifts in amino-acid preferences, we next looked at sites with other special structural or functional properties. One group of functionally important sites are those that comprise the RNA-binding groove of NP. These RNA-binding sites have low *RMSD_within_* (Figure 4A and B), indicating below-average noise among replicates. RNA-binding sites also have low *RMSD_corrected_* (Figure 5B, Figure 6A). These results are consistent with the expectation that RNA-binding sites in NP are under strong and conserved functional constraint, since RNA binding is essential for viral genome packing, transcription, and replication.

**Figure 6:**
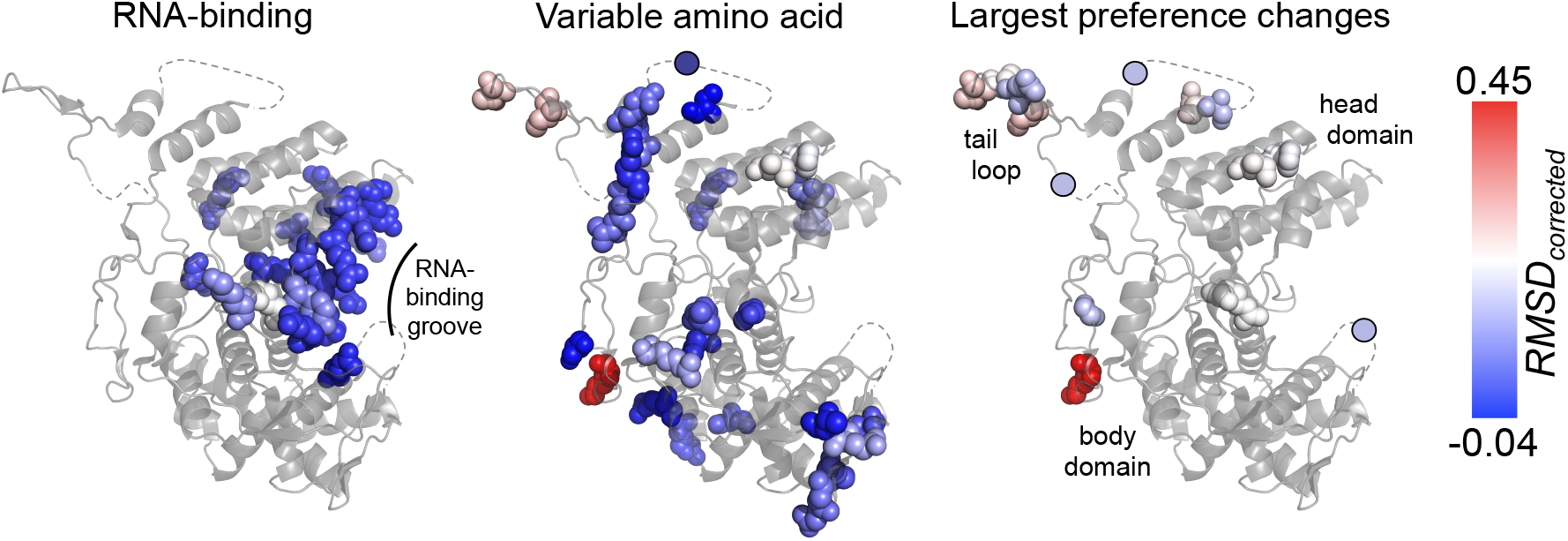
Magnitude of the shift in amino-acid preferences mapped on the NP structure. *RMSD_corrected_* values for each site are used to color space-filling models for the indicated sites in the NP crystal structure (PDB ID 2IQH, chain C; Ye et al., 2006)). Sites are shown as circles when in regions that are not present in the crystal structure (dashed lines). Blue represents small shifts in amino-acid preferences between PR/1934 and Aichi/1968; red represents large shifts. *Variable amino acid* refers to sites where the wild-type residue differs between PR/1934 and Aichi/1968 NP. *Largest preference changes* refers to sites where the null hypothesis is rejected using exact randomization testing with a false discovery rate of 5%.

We next hypothesized that sites in structural proximity to evolutionarily variable sites may experience shifts in amino-acid preferences due to changes in the surrounding biochemical environment. We identified sites directly contacting the evolutionarily variable residues, and found that they do not have *RMSD_corrected_* values that differ from other sites (Figure 5B). Therefore, we are unable to identify any preferential tendency for substitutions to drive shifts in amino-acid preference at other sites in direct contact with the substituted residue.

The 14 sites with the clearest shifts in amino-acid preferences are distributed throughout the surface of NP in the body, head, and tail loop domains (Figure 6). Six of the 14 sites are located in the flexible tail loop, which inserts into a neighboring monomer during NP oligimerization. This suggestive clustering led us to test whether there was a significant tendency for the 14 sites with clearest shifts in preferences to be spatially clustered in NP’s structure. We calculated the distance between sites as the minimum distance between side chain atoms (using the alpha carbon for glycine). Eleven of the 14 sites with clearest shifts in preferences are resolved in the crystal structure, and of these 11 sites the median distance to the nearest neighbor among the 10 remaining sites is 5.8 Ångströms, which is significantly less than expected by chance for random selections of 11 sites (10.8 Ångströms, p=0.028). Thus, the clearest shifts in preferences between these two homologs occur in small clusters of proximal sites more often than in single isolated sites. This pattern also holds when considering the 76 sites identified by the simulation null distribution: among the 66 that are resolved in the crystal structure the median distance to the nearest neighbor is 4.5 Ångströms compared to a median distance to nearest neighbor among random selections of 66 sites of 5.0 Ångströms (p = 0.021). Therefore, sites with shifted preferences appear to cluster in NP’s structure, even if they are not usually in direct physical contact with variable residues.

Overall, these results indicate that sites with evolutionarily variable amino-acid identity are more likely than conserved sites to exhibit shifts in amino-acid preferences, and that sites with shifted preferences tend to cluster in NP’s structure. However, the majority of sites with variable identity do not exhibit large shifts in amino-acid preference, and overall, only between 3% and 15% (depending on the method used to generate the null distribution) of sites in NP undergo shifts in amino-acid preferences that are sufficiently large to justify rejecting the null hypothesis that the preferences are identical between homologs. Importantly, statistical significance does not necessarily imply a large magnitude in effect size – and indeed, with just a handful of exceptions (most prominently site 470), even the shifted sites are vastly more similar in their preferences than typical pairs of sites in non-homologous proteins.

### Experimentally informed site-specific substitution models describe vast swaths of nucleoprotein evolution

We next quantitatively assessed how well our experimentally measured amino-acid preferences reflected the actual constraints on NP evolution. To do so, we used the amino-acid preferences to inform sitespecific phylogenetic substitution models. We have previously shown that substitution models informed by experimentally measured site-specific amino-acid preferences greatly outperform common non-site-specific codon-substitution models (Bloom, 2014a,b; Thyagarajan and Bloom, 2014).

In the prior work, site-specific amino-acid preferences were experimentally measured in a single sequence context. Here, we asked whether combining the preferences measured in the two different sequence contexts of Aichi/1968 and PR/1934 would more accurately describe NP sequence evolution. Any improvement could be due to two effects: First, a combined substitution model might better reflect the evolutionary average of the amino-acid preferences at sites with significant changes in preferences over time. Second, combining data from multiple experiments should reduce noise and yield more accurate site-specific amino-acid preferences.

#### Combining deep mutational scanning datasets from nucleoprotein homologs improves phylogenetic fit

To compare the performance of different substitution models, we used a likelihood-based framework. We first built a maximum-likelihood tree for NP sequences from human influenza using `CodonPhyML` (Gil et al., 2013) with the codon-substitution model of Goldman and Yang (1994) (GY94) (Figure 1). We fixed this tree topology and used `HyPhy` to optimize branch lengths and model parameters for each substitution model by maximum likelihood. The relative fits of the substitution models were evaluated using the Akaike information criterion (AIC) (Posada and Buckley, 2004).

We tested experimentally informed substitution models derived from the Aichi/1968 and PR/1934 mutational scans either alone or in combination. The Aichi/1968 model used amino-acid preferences averaged across the *current study* and the *previous study*. To build a combined substitution model based on both NP homologs, we averaged the amino-acid preferences for the Aichi/1968 and PR/1934 homologs (Aichi/1968 + PR/1934). Each substitution model had five free parameters that were fit by maximum likelihood: four nucleotide mutation rates and a stringency parameter *β* that accounts for the possibility of a different strength of selection in natural sequence evolution compared to the mutational-scanning experiments (Bloom, 2014b). Importantly, the amino-acid preferences themselves are not free parameters, as they are independently measured by experiments that do not utilize information from the naturally occurring NP sequences.

As a comparison to the experimentally informed substitution models, we also tested the non-site-specific GY94 model. Relative to the experimentally informed substitution models, the GY94 model includes more free parameters including equilibrium codon frequencies, a transition-transversion ratio, and parameters describing gamma distributions of the nonsynonymous-synonymous ratio and substitution rate across sites (Yang, 1994; Yang et al., 2000).

The Aichi/1968 and PR/1934 experimentally informed models described the human NP phylogeny far better than the non-site-specific GY94 model (Table 1). Strikingly, combining amino-acid preferences from both NP homologs (Aichi/1968 + PR/1934) resulted in a greatly improved substitution model (Table 1). For each experimentally informed model, the stringency parameter *β* fit with value greater than 1 (average *β* = 2.5), consistent with the idea that selection during natural evolution is more stringent than our laboratory selection.

**Table 1:**
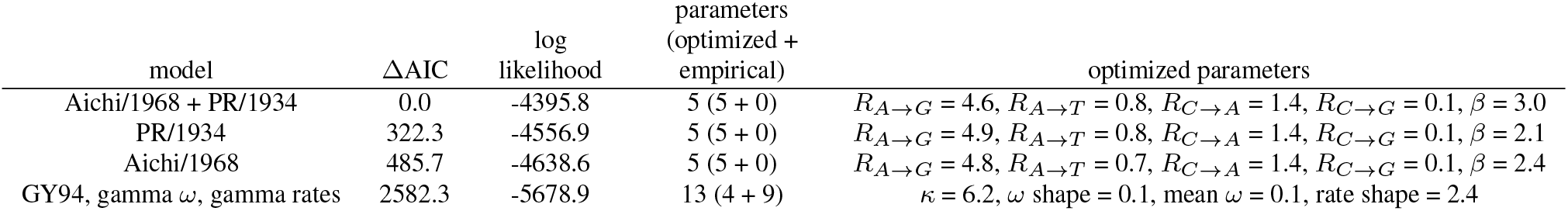
Combining experimental data improves phylogenetic fit to NPs from human influenza. Substitution models are sorted by ΔAIC, and the corresponding log likelihoods, number of free parameters, and values of optimized parameters are shown. Log likelihoods for each model were calculated through maximum-likelihood optimization of branch lengths and model parameters given the fixed tree topology of human NPs shown with blue lines in Figure 1. The only parameters in the experimentally informed models are the four nucleotide mutation rates and the stringency parameter *β*. The non-site-specific GY94 model (Goldman and Yang, 1994) has nine empirical nucleotide equilibrium frequencies (Pond et al., 2010), and optimized parameters describing the transition-transversion ratio (*?*), the gamma distribution of the nonsynonymous-synonymous ratio (*ω*) (Yang et al., 2000), and the gamma distribution of substitution rates (Yang, 1994). In the Aichi/1968 model, the preferences from *current study* and *previous study* have been averaged.

#### Experimentally informed models also describe the evolution of more diverged non-human influenza strains

Given the success of the experimentally informed substitution models in describing the human NP phylogeny, we asked whether these models could be extended to more diverged NPs from non-human influenza strains. We expect these models to exhibit good fit if the NP site-specific aminoacid preferences are mostly conserved across these viral strains. We examined NPs from influenza strains from three hosts: swine, equine, and avian. The average protein-sequence identity between human NPs and swine, equine, and avian NPs was 91%, 91%, and 93% respectively.

We built a phylogenetic tree of NPs of influenza viruses from human, swine, equine, and avian hosts (Figure 1, Supplementary file 4). As previously reported, the avian sequences could be divided into western and eastern hemispheric clades, and the swine sequences consisted of the North American Classical H1N1 clade and the more recent Eurasian H1N1 clade (Worobey et al., 2014). Using this tree, we performed a phylogenetic analysis similar to that described above for human influenza NPs.

Again, the experimentally informed models greatly outperformed the non-site-specific GY94 model, and combining the Aichi/1968 and PR/1934 models resulted in a far superior model (Table 2). Since the amino-acid preferences were experimentally measured for human NP, we wanted to ensure that this superior performance was not driven solely by the human clade of the tree. We separately fit subtrees consisting only of swine, equine, or avian NP sequences (Supplementary table 1, Supplementary table 2, Supplementary table 3). Each subtree showed the same trend as the full tree: the experimentally informed models were superior to the GY94 model, and combining data from the two NP homologs resulted in large improvements in likelihood. Therefore, site-specific amino-acid preferences of NP are sufficiently conserved across influenza A lineages that substitution models informed by deep mutational scanning of human influenza NP homologs can be extended to the NPs of influenza from other hosts.

**Table 2:**
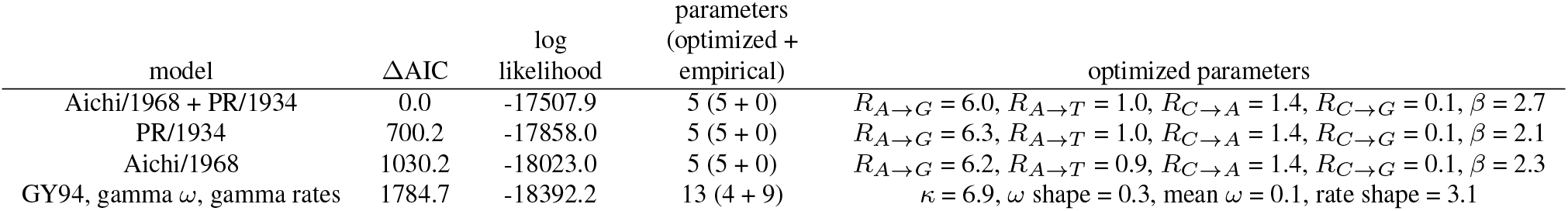
Combining experimental data improves phylogenetic fit to NPs from human, swine, equine, and avian influenza. This table differs from Table 1 in that it fits the combined tree of human, swine, equine, and avian NPs in Figure 1.

| model | $\Delta AIC$ | log<br>likelihood | parameters<br>(optimized +<br>empirical) | optimized parameters |
| --- | --- | --- | --- | --- |
| Aichi/1968 + PR/1934 | 0.0 | -17507.9 | 5 (5 + 0) | $R_{A \rightarrow G} = 6.0, R_{A \rightarrow T} = 1.0, R_{C \rightarrow A} = 1.4, R_{C \rightarrow G} = 0.1, \beta = 2.7$ |
| PR/1934 | 700.2 | -17858.0 | 5 (5 + 0) | $R_{A \rightarrow G} = 6.3, R_{A \rightarrow T} = 1.0, R_{C \rightarrow A} = 1.4, R_{C \rightarrow G} = 0.1, \beta = 2.1$ |
| Aichi/1968 | 1030.2 | -18023.0 | 5 (5 + 0) | $R_{A \rightarrow G} = 6.2, R_{A \rightarrow T} = 0.9, R_{C \rightarrow A} = 1.4, R_{C \rightarrow G} = 0.1, \beta = 2.3$ |
| GY94, gamma $\omega$, gamma rates | 1784.7 | -18392.2 | 13 (4 + 9) | $\kappa = 6.9, \omega \text{ shape} = 0.3, \text{mean } \omega = 0.1, \text{rate shape} = 3.1$ |

#### Combining data from NP homologs improves phylogenetic fit to sites with shifted preferences

The results above show that the experimentally informed substitution models improved phylogenetic fit relative to the non-site-specific model, and that combining data from two NP homologs resulted in the best model. This increased performance when combining data may come from more accurate measurement of amino-acid preferences due to more replicates, or from averaging amino-acid preferences over multiple sequence contexts. To examine these possible explanations, we analyzed which sites in NP were more accurately modeled when the Aichi/1968 and PR/1934 experimental models were combined. This analysis was performed using the full phylogenetic tree of NP sequences (Figure 1).

While fixing the branch lengths and model parameters to their maximum-likelihood values for each model, we calculated for each site the difference in likelihoods (Δlog-likelihood) when the site was modeled using the combined Aichi/1968 + PR/1934 model compared to using the Aichi/1968 model. We binned sites into quintiles of Δlog-likelihood. Sites in the top quintile had the greatest increases in likelihood when the Aichi/1968 and PR/1934 models were combined. Overall 67% of sites in NP had increased likelihoods under the Aichi/1968 + PR/1934 model.

To determine whether these improved likelihoods came from lower noise in the combined experimental model, we used the *RMSD_within_* statistic. Sites with greater variance in amino-acid preferences across experimental replicates have higher *RMSD_within_* scores. We analyzed the distribution of the *RMSD_within_* scores for sites within each quintile (Figure 7). The top and bottom quintiles did not have significantly different *RMSD_within_* distributions, indicating that sites prone to experimental noise contributed both positively and negatively to the tree likelihood when experimental datasets were combined. Thus, the improved modeling with the combined dataset was not chiefly due to reduced experimental noise.

**Figure 7:**
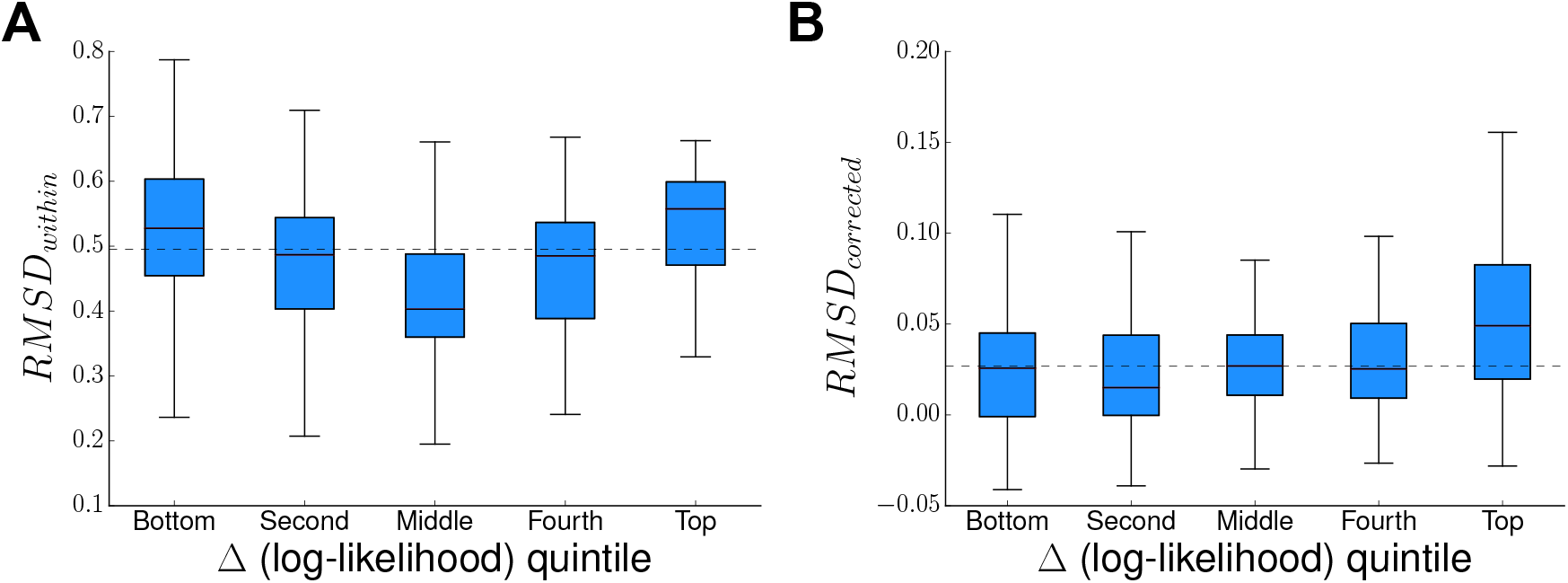
NP sites that are better described by combining data from both homologs have shifted amino-acid preferences. The change in per-site likelihood in going from the Aichi/1968 model to the Aichi/1968 + PR/1934 model was plotted against the per-site *RMSD_within_* (A) or persite *RMSD_corrected_* (B). Sites were ranked by Δ(log-likelihood), divided into quintiles, and the persite *RMSD_within_* or per-site *RMSD_corrected_* for sites in each quintile was displayed as a box and whisker plot. Outlier sites beyond the interquartile range are omitted. Quintiles are ordered left to right from least improved likelihoods to most improved likelihoods under the combined model. The median *RMSD_within_* or *RMSD_corrected_* is shown as a horizontal, dashed line. Sites with the most improved likelihoods did not have significantly higher variation in amino-acid preferences (high *RMSD_within_*) across replicate measurements on the same homolog. However, these sites did have significantly higher differences in amino-acid preferences between Aichi/1968 and PR/1934 (high *RMSD_corrected_*).

Next, to determine whether the improved likelihoods were driven by sites with different preferences between the two NP homologs, we used the *RMSD_corrected_* statistic (Figure 7). If the improvements under the combined model came from sites with different amino-acid preferences between Aichi/1968 and PR/1934, then we would expect that the sites with the greatest increases in likelihood would also have the greatest *RMSD_corrected_* values. This was indeed the case, as sites in the top quintile of loglikelihoods had the highest median *RMSD_corrected_*. The *RMSD_corrected_* scores in the top quintile were significantly different from those in the lower quintiles (Mann-Whitney U with Bonferroni correction *p<* 0.002), whereas there were no significant differences in the *RMSD_corrected_* scores when comparing the lower quintiles. Therefore, improvements in the combined model were partly due to better describing those sites that had the largest shifts in amino-acid preferences over evolutionary time.

## Discussion

Determining the extent to which site-specific amino-acid preferences shift during evolution is important for evaluating how well experimental measurements can be extrapolated across homologs, and for guiding the development of site-specific phylogenetic substitution models. We have performed the first comprehensive assessment of the conservation of site-specific amino-acid preferences by using deep mutational scanning to measure the effects of all mutations on two closely related homologs of influenza NP.

We found that for the majority of sites, any shift in amino-acid preferences between homologs was smaller than the noise in our experiments. We could reject the null hypothesis that the amino-acid preferences were identical among homologs for only between 3% and 15% of all sites, depending on the method used to generate the null distribution. Furthermore, even for those sites for which we could reject the null hypothesis of identical preferences between homologs, the magnitude of shifts tended to be small. Only a handful of the 497 sites exhibited shifts in preference between homologs with a magnitude comparable to the average difference between sites in non-homologous proteins. Sites that varied in amino-acid identity between the two homologs were more likely to have a detectable shift in amino-acid preferences – but even among variable sites, there was usually no shift. Admittedly, our experiments had substantial noise, so it is likely that other sites have undergone subtle shifts below our limit of detection. However, the fact that the preferences for the two NP variants are strongly correlated with each other but completely uncorrelated with those for the non-homologous HA shows that the site-specific amino-acid preferences of homologs are tremendously more similar than those of unrelated proteins.

This general conservation of site-specific amino-acid preferences does not imply an absence of epistasis during NP’s evolution. For instance, our results show that some (as yet mechanistically uncharacterized) epistatic interaction with other sites has driven a strong shift in the amino-acid preferences at site 470. At other sites, smaller shifts in amino-acid preferences are still certain to induce evolutionarily important epistasis, since natural selection is highly discerning. Indeed, we have previously demonstrated epistasis among mutations to NP (Gong et al., 2013), indicating that NP is no different than the many other proteins for which evolutionarily relevant epistasis has been identified (Weinreich et al., 2006; Ortlund et al., 2007; da Silva et al., 2010; Lunzer et al., 2010; Natarajan et al., 2013; Podgornaia and Laub, 2015). Our key result is not that epistasis is absent, but rather that its frequency and magnitude are sufficiently low that the amino-acid preferences for most sites are are still vastly more similar between homologs than between non-homologous proteins.

The implications of this finding are illustrated by the second part of our study, which shows that the experimentally measured site-specific amino-acid preferences can inform phylogenetic substitution models that greatly outperform non-site-specific models even for more diverged NP homologs. It is well known that the actual constraints on protein evolution involve cooperative interactions among sites (Zuckerkandl and Pauling, 1965; DePristo et al., 2005; Harms and Thornton, 2013), and so substitution models that treat sites either independently or identically are obviously imperfect. But computational biology must balance realism with tractability. Site-independent but site-specific substitution models are becoming feasible for real-world datasets (Lartillot and Philippe, 2004; Le et al., 2008; Wang et al., 2008; Rodrigue et al., 2010; Bloom, 2014a,b), but approaches that relax the assumption of independence among sites remain in their infancy (Choi et al., 2007; Bordner and Mittelmann, 2014). Are amino-acid preferences sufficiently conserved for site-independent but site-specific models to represent substantial improvements over existing non-site-specific alternatives? Both our experimental and computational results answer this question with a resounding yes.

Why are the site-specific amino-acid preferences mostly conserved? As is the case for virtually all proteins (Chothia and Lesk, 1986; Sander and Schneider, 1991), the structure of NP is highly conserved among homologs (Ye et al., 2006; Das et al., 2010), and sites in specific structural contexts often have propensities for certain amino acids (Chou and Fasman, 1974; Richardson and Richardson, 1988; Lim and Sauer, 1991). In addition, selection for protein stability is a major constraint on evolution (DePristo et al., 2005; Bloom et al., 2005), and experiments on NP (Ashenberg et al., 2013) and other proteins (Risso et al., 2015; Serrano et al., 1993) have shown that the effects of mutations on stability are similar among homologs. Therefore, conserved structural and stability constraints probably naturally lead to substantial conservation of site-specific amino-acid preferences. We refer the reader to an excellent recent study by Risso et al. (2015) for a more biophysically nuanced discussion of these issues.

The extent to which site-specific amino-acid preferences will be conserved among more distant homologs remains an open question. Computational simulations of the divergence of distant homologs have been used to argue that preferences shift substantially (Pollock et al., 2012), but the reliability of such simulations is unclear since computational predictions of the effects of even single amino-acid mutations are only modestly accurate (Kellogg et al., 2011; Potapov et al., 2009). The only direct experimental data come from a study showing that the effects of a handful of mutations on stability are mostly conserved among homologs with about 50% protein-sequence identity (Risso et al., 2015). More comprehensive determination of the relationship between sequence divergence and shifts in site-specific amino-acid preferences therefore remains an important topic for future work.

## Methods

### Availability of data and computer code

FASTQ files can be accessed at the Sequence Read Archive (SRA Accession SRP056028). The computer code necessary to reproduce all the analysis in this work is available at https://github.com/mbdoud/Compare-NP-Preferences.

### Deep mutational scanning of two influenza nucleoprotein homologs

We performed deep mutational scanning of influenza nucleoprotein (NP) in three biological replicates for A/PR/1934 (H1N1) and two biological replicates for A/Aichi/1968 (H3N2) (termed here as Aichi/1968 *current study*). We broadly followed the methods used for mutagenesis, viral rescue, deep sequencing, and inference of amino-acid preferences from sequence data described in (Bloom, 2014a), with the following notable changes to the protocol.

#### Codon mutagenesis

For each replicate mutant library, we followed the mutagenesis protocol as previously described (Bloom, 2014a), but performed two rounds of mutagenesis instead of three to decrease the average number of mutations per clone. After ligation of mutagenized PCR products to the pHW2000 (Hoffmann et al., 2000) plasmid backbone, multiple parallel transformations and platings were combined to ensure that each replicate library contained more than 10^6^ unique transformants. Sanger sequencing of 30 clones from each homolog revealed that the number of mutations per clone was approximately Poisson distributed with an average of 1.7 mutations per clone for the PR/1934 libraries and 2.1 mutations per clone for the Aichi/1968 libraries, with mutations distributed uniformly across the length of the gene.

#### Growth of mutant virus libraries

We used reverse genetics (Hoffmann et al., 2000) to rescue viruses carrying mutant NP genes. Co-cultures of 293T and MDCK-SIAT1 cells were plated 16 hours prior to transfection in D10 media (DMEM supplemented with 10% FBS, 100 U/mL of penicillin,100 *μ*g/mL of streptomycin, and 2 mM L-glutamine) at cell densities of 3x10^5^ 293T/mL and 2.5x10^4^ MDCK-SIAT1/mL. Co-cultures were transfected using BioT transfection reagent (Bioland Scientific) with a mixture of 250 ng of each of the eight reverse genetics plasmids per well in 6-well plates. In order to circumvent the possibility of rare mutants with exceptional replication fitness growing to high frequencies and limiting the growth of other mutants, we divided each transfection into multiple tissue-culture wells.

For the PR/1934 libraries, we rescued viruses containing the mutagenized PR/1934 NP with the seven remaining PR/1934 viral gene segments, and each replicate mutant library was transfected into the twelve wells of two 6-well plates. For the Aichi/1968 libraries, we used a viral rescue protocol that increases the number of parallel transfections and uses 293T cells that constitutively express protein V from hPIV2. This protein targets STATI for degradation, thereby inhibiting type I interferon signaling (Andrejeva et al., 2002). We rescued these Aichi/1968 virus libraries by transfecting the Aichi/1968 NP mutant library along with PB1/PB2/PA from Nanchang/933/1995 (using the plasmids in (Gong et al., 2013) and HA/NA/M/NS from WSN/1933 into 48 wells of eight 6-well plates. For both homologs, in parallel, we performed similar transfections using the corresponding unmutated NP genes to grow unmutated virus.

At 24 hours after transfection, co-culture media was aspirated, cells were rinsed with PBS, and the media was changed to influenza growth media (OptiMEM I media (Gibco) supplemented with 0.01% FBS, 0.3% BSA, 100 U/mL of penicillin,100 *μ*g/mL of streptomycin, 100 *μ*g/mL calcium chloride, and 3 *μ*g/mL TPCK-trypsin). Co-culture supernatant was collected 72 hours after transfection, clarified by centrifugation at 2,000x*g* for 5 min, aliquoted and stored at -80*°* C.

Since many of the virions obtained from transfection with mutant NP library plasmids are likely to have originated in cells that contained more than one mutant NP gene and therefore might carry NP genes and NP proteins with different mutations, we passaged viruses in MDCK-SIAT1 cells at a low multiplicity of infection (MOI) to enforce genotype-phenotype linkage. We titered viruses from thawed transfection supernatant aliquots for each replicate virus library using the TCID50 protocol described in (Thyagarajan and Bloom, 2014). We then passaged viral libraries in MDCK-SIAT1 cells. Cells were plated in D10 media at 2x10^5^ cells/mL. After 16 hours, the media was changed to influenza growth media containing diluted transfection supernatant virus. PR/1934 libraries were each passaged in 20 wells of 6-well dishes at an MOI of 0.05 TCID50/cell, and Aichi/1968 libraries were each passaged in eight 10-cm dishes at an MOI of 0.1 TCID50/cell. After 48 hours, supernatant was clarified by centrifugation at 2,000x*g* for 5 min, aliquoted and stored at -80*°* C.

#### Sample preparation and deep sequencing

For each virus sample to be sequenced, 10 mL of clarified viral passage supernatant was centrifuged at 64,000x*g* for 1.5 hours to pellet viruses. RNA was extracted using the Qiagen RNEasy kit by lysing viral pellets in buffer RLT and following the manufacturer’s recommended protocol. The NP gene was reverse transcribed using AccuScript High-Fidelity Reverse Transcriptase (Agilent Technologies) from both positive-sense and negative-sense viral RNA templates using the primers PR8-NP-RT-F (5’-agcaaaagcagggtagataatcactcactgagtgac-3’) and PR8-NP-RT-R (5’-agtagaaacaagggtatttttcttta-3’) for PR/1934 viruses or the primers 5’-BsmBI-Aichi68-NP (5’-catgatcgtctcagggagcaaaagcagggtagataatcactcacag-3’) and 3’-BsmBI-Aichi68-NP (5’-catgatcgtctcgtattagtagaaacaagggtatttttcttta-3’) for Aichi/1968 viruses.

To ensure a sufficiently large number of unique RNA molecules were reverse transcribed, we used qPCR (SYBR Green Real-Time PCR Master Mix, Life Technologies) using primers qWSN-NP-for (5’-ACGGCTGGTCTGACTCACAT-3’) and qPR8-NP-rev (5’-TCCATTCCGGTGCGAACAAG-3’) to quantify the concentration of first-strand cDNA molecules against a standard curve of linear NP amplicons quantified by Quant-iT PicoGreen dsDNA Assay Kit (Life Technologies). We then made PCR amplicons with KOD DNA Polymerase (Merck Millipore) using at least 1x10^9^ first-strand cDNA molecules as template in each reaction for viral gene sequencing. We also made PCR amplicons using 10 ng of the indicated plasmids for plasmid sequencing. For each biological replicate, we generated these PCR amplicons with 25 cycles of amplification using unmutated NP plasmid, mutated NP plasmid, NP cDNA from unmutated virus, and NP cDNA from mutated virus as template for the **DNA**, **mutDNA**, **virus**, and **mutvirus** samples, respectively.

To reduce the sequencing error rate, we developed a sequencing sample preparation protocol that results in sequencing libraries with inserts approximately 150 bp long. This allowed us to use paired-end 150 bp sequencing to achieve mostly overlapping reads so that sequencing errors resulting in mismatches between the two reads could be identified and ignored during data analysis. To make these sequencing libraries, we gel-purified the **DNA**, **mutDNA**, **virus**, and **mutvirus** PCR amplicons and sheared 1 *μ*g of each amplicon using Covaris to a median size of approximately 150 bp. We followed the modified Illumina paired-end library preparation protocol provided in (Henikoff et al., 2011) for end repair, 3’ A overhang, and adapter ligation steps, using Zymo DNA Clean & Concentrator columns (Zymo Research) or Ampure XP (Beckman Coulter) magnetic beads for DNA clean-up after shearing, end repair, and 3’ A overhang steps. Barcoded Y-adapters were made by annealing 10 *μ*L of 100 *μ*M PAGE purified universal adapter (5’-AATGATACGGCGACCACCGAGATCTACACTCTTTCCCTACACGACGCTC TTCCGATC*T- 3’, where * indicates phosphorothioate bond) to 10 *μ*L of 100 *μ*M PAGE purified barcoded adapter (5’-PGATCGGAAGAGCACACGTCTGAACTC CAGTCACNNNNNNATCTCGTATGCCGTCTTCTGCTT*G-3’, where P indicates 5’ phosphorylation, * indicates phosphorothioate bond, and NNNNNN indicates sample-specific barcode). Each 20 *μ*L mixture (one mixture for each barcode sequence) was annealed by heating to 95*°* C for 5 minutes and cooling at 0.3*°* C/second to 4*°* C. The resulting Y-adapters were diluted to 25 *μ*M by adding 20*μ*L 10 mM Tris pH 7.5 and stored in 4 *μ*L aliquots at -20*°* C. Y-adapters with unique barcodes (ATCACG, ACTTGA, TAGCTT, GGCTAC, TTAGGC, GATCAG, ACTGAT, CGTACG, CGATGT, TGACCA, CAGATC, and CCGTCC) were ligated to samples derived from each biological replicate of each amplicon and ligation products were purified using 0.8X bead-to-sample ratio Ampure XP.

Purified adapter-ligated products for each sample were quantified by Quant-iT PicoGreen dsDNA Assay Kit (Life Technologies) and 25 ng was used as template for a 4-cycle PCR using Phusion High-Fidelity Polymerase (Thermo Scientific) to amplify inserts with adapters properly ligated on both sides. This amplification step was performed with the following components: 25 ng template DNA, 5 *μ*L 5X Phusion buffer, 2.5 *μ*L mixture of each dNTP at 2.5 mM, 2 *μ*L forward primer at 10 *μ*M (5’-AATGATACGGCGACCACCGAGATCTACACTCTTTCCCTACACGA-3’), 2 *μ*L reverse primer at 10 *μ*M (5’-CAAGCAGAAGACGGCATACGAGAT-3’), and 0.25 *μ*L Phusion polymerase in a final reaction volume of 25 *μ*L. PCR products were purified using 1.0X bead-to-sample ratio Ampure XP and quantified using PicoGreen. Samples were pooled in equal amounts and size-selected on a 2.0% agarose gel for fragments between 240 bp and 300 bp, which contain sequencing inserts in the size range of 120-180 bp. The size-selected sample was then sequenced at the Fred Hutchinson Genomics Core on an Illumina HiSeq 2500 using a paired-end 150 bp sequencing strategy in rapid run mode.

#### Analysis of deep sequencing data

Sequencing data processing was performed using the software package mapmuts (Bloom, 2014a). Briefly, for each replicate sample of **DNA**, **mutDNA**, **virus**, and **mutvirus**, paired reads were stripped of any adapter sequence and aligned to each other. Read pairs were discarded if any of the following criteria were met: less than 100 bp of overlap between reads, average Q-score less than 25 across either read, more than 5 ambiguous nucleotides (N nucleotides) in either read, or more than 1 mismatch in the overlap between reads. Retained read pairs were then aligned to the appropriate reference NP gene sequence for PR/1934 or Aichi/1968 NP, and read pairs with more than 10 mismatches to the reference sequence or with any gaps or insertions were discarded. Once aligned to the reference sequence, codon identities at every position were called only if all three nucleotides in the codon matched unambiguously in both reads. The total number of codon identities at every codon position in the coding region were totaled for each sample (**DNA**, **mutDNA**, **virus**, and **mutvirus**), separately for each biological replicate.

#### Inference of amino-acid preferences

We specify that at every site *r* in the protein, there is an inherent preference *π*_*r,a*_ for every amino acid *a*, and we specify that Σ_*a*_ *π*_*r,a*_ = 1. The preference *π*_*r,a*_ can be considered to be the expected frequency of amino acid *a* at site *r* in a mutant virus library after viral growth from a starting plasmid mutant library that contains equal numbers of every amino acid encoded at site *r*. Thus, mutations to amino acids with high preferences are beneficial and will be selected for during viral growth, and mutations to amino acids with low preferences will inhibit viral growth and will be selected against. Since the plasmid mutant libraries we generated contain on average more than one mutation per clone, the amino-acid preferences we measure represent an average preference in a variety of genetic backgrounds very similar to the starting sequence.

Let *A*(*x*) represent the amino acid encoded by codon *x* and let *C* represent the set of all codons. The effect of the preference *π*_*r,A*(*x*)_ on the frequency *f* of observing codon *x* at site *r* in the mutant virus library sample **mutvirus** is given by:

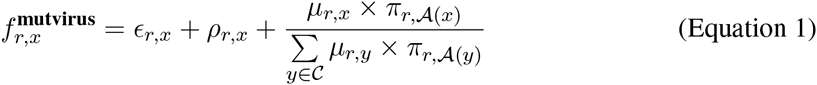

where *∈*_*r,x*_ is the rate of PCR and sequencing errors at site *r* resulting in codon *x*, *ρ*_*r,x*_ is the rate of reverse transcription errors at site *r* resulting in codon *x*, and *μ*_*r,x*_ is the frequency of codon *x* at site *r* in the plasmid mutant library **mutDNA**.

We inferred the amino-acid preferences independently for each biological replicate using the Bayesian algorithm described in (Bloom, 2015) as implemented in `dms_tools` where codon counts in the **DNA**, **virus**, and **mutDNA** samples are used to infer the unknown parameters *E*, *ρ*, and *μ* at each site.

Amino-acid preferences for Aichi/1968 NP were previously published in (Bloom, 2014a), where 8 biological replicates of the entire experiment were performed. In this work we report two additional biological replicates of the deep mutational scanning experiment for Aichi/1968. We will distinguish the two data sets when they are used separately for comparison as Aichi/1968 *previous study* and Aichi/1968 *current study*, and we will call the combined dataset of all 10 biological replicates for this homolog *Aichi/1968*.

### Comparison of site-specific amino-acid preferences between homologs

#### Quantifying the magnitude of amino-acid preference difference between homologs

At every site in the protein, each replicate deep mutational scanning experiment allows for the inference of an aminoacid preference distribution 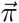 that provides the preference at that site for all 20 amino acids. We used the Jensen-Shannon distance metric (the square root of the Jensen-Shannon divergence) to quantify the distance *d* between two amino-acid preference distributions:

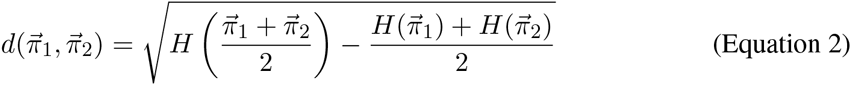

where 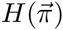 is the Shannon entropy of the amino-acid preference distribution 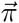. The Jensen-Shannon distance metric quantifies the similarity between two amino-acid preference distributions, ranging from 0 (identical distributions) to 1 (completely dissimilar distributions). The average distance *d* between amino-acid preferences inferred from replicate experiments in the same homolog varies across sites. In other words, at some sites in the protein 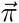 is measured with greater precision than others. We therefore sought to develop, for every site *r*, a quantitative measure of the magnitude of change in 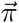 between homologs that corrects for the variation in 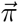 within replicate experiments of the same homolog.

For two groups of replicate mutational-scanning experiments *A* and *B* done in different homologs, each containing several replicate inferences of 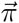 for every site, we calculate the root-mean-square distance at site *r* over all pairwise comparisons of 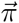 measured in replicate experiments *i* (from group *A*) and *j* (from group *B*):

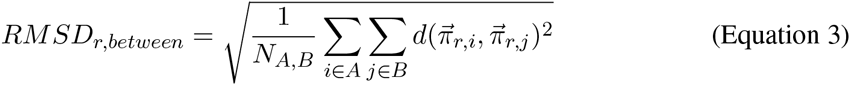

where *N*_*A,B*_ is the total number of non-redundant pairwise comparisons between replicate preferences measured from groups A and B. At the same site, to estimate the amount of experimental noise within replicates of the same homolog, we calculate the root-mean-square distance over all pairwise comparisons of 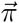 *within* the same group of replicate experiments, and average this site-specific noise estimate across the two groups:

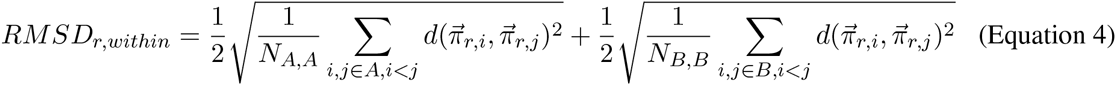

where *N*_*A,A*_ and *N*_*B,B*_ are the number of non-redundant pairwise comparisons between replicates within groups A and B, respectively. We then subtract the magnitude of the noise at this site observed *within* groups from our measurement of the difference in amino-acid preferences seen *between* groups to obtain a corrected value for the change in 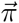 at site *r* between homologs:

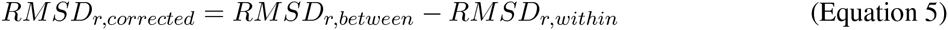

It is possible that the observed variation within groups is greater than the observed variation between groups, resulting in negative *RMSD_corrected_*.

#### Identifying sites with statistically significant changes in amino-acid preference

To determine whether site-specific *RMSD_corrected_* values are significantly larger than expected if amino-acid preferences are unchanged between homologs, we applied two methods to generate null distributions of *RMSD_corrected_* values. First, we used exact randomization testing to make all possible shuffles of the replicate homolog datasets into the two groups *A* and *B*. For each permutation, we calculated the *RMSD_corrected_* at every site, and the results are combined for all permutations. If there are no differences in preferences between homologs, the distribution of scores generated through randomization should be similar to the distribution of scores from the actual experiment.

We next observed that the overall correlation of amino-acid preferences across all sites between replicates can vary between experiments. For instance, the average Pearson’s correlation between PR/1934 replicates is 0.59, the correlation between Aichi/1968 replicates in the *previous study* is 0.50, and the correlation between Aichi/1968 replicates in the *current study* is 0.74. We considered whether the varying precision between homologs might lead to biases in the calculated *RMSD_corrected_*.

To test this, we generated a second null distribution of *RMSD_corrected_* under the hypothesis that the “true” amino-acid preferences are the same for both homologs and can be approximated by averaging the mean observed preferences for each homolog:

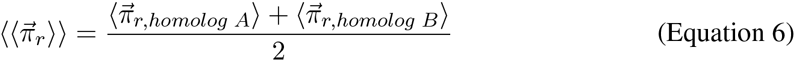

Under this hypothesis, the observed differences in amino-acid preferences between homologs is solely due to the different amounts of experimental noise between replicates of each homolog. To model the effects of this noise on our analysis, we drew replicate simulated amino-acid preferences at each site *r* from a Dirichlet distribution with mean centered on the “true” amino-acid preferences:

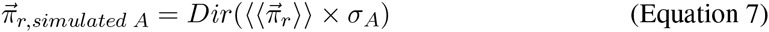

where *σ*_*A*_ is a scaling factor that is chosen to yield simulated replicate preferences across the entire protein that have an average Pearson’s correlation between replicates equal to the correlation between experimental replicates. In other words, we simulate replicate amino-acid preference measurements with noise tuned to match the actual noise in each experiment. For each simulated experiment, we simulated the same number of replicates that were performed experimentally, and calculated *RMSD_corrected_* for all sites. We ran the entire simulation 1000 times, combining all *RMSD_corrected_* values to obtain a null distribution.

We then separately used the two null distributions (generated through randomization or simulation) to assign p-values to site-specific *RMSD_corrected_* at each site *r*:

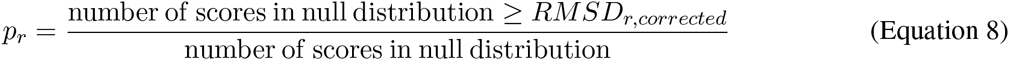

To control the false discovery rate across the 497 sites tested for significance, we used the procedure of Benjamini and Hochberg (1995).

#### Structural analysis of sites with preference changes

We used the crystal structure of the influenza A H1N1 WSN/1933 NP (PDB ID 2IQH, chain C; Ye et al., 2006) to calculate distances between sites. Distances between sites were defined as the minimum distance between any side chain atoms distal to the alpha carbons of each site (the alpha carbon was used for all glycine residues). A distance cutoff of 4.5 Ångströms was used to define sites that are in contact with evolutionarily variable sites. To test for spatial clustering of a group of *N* sites, the distribution of *N* distances to the nearest neighbor of the remaining *N -* 1 sites was compared to a null distribution of distances calculated the same way for 1000 random selections of sites of size *N*. One-sided P-values were computed using the Mann-Whitney U test.

### Phylogenetic analysis

#### Experimental substitution model overview

We used a previously described approach to build sitespecific substitution models for influenza nucleoprotein (Bloom, 2014a,b). Briefly, this approach calculates the codon-substitution rate at each site in nucleoprotein based on the rate at which nucleotide mutations arise and the level of selection acting on these new mutations. The rate of codon substitution, *P*_*r,xy*_, at site *r* of codon *x* to a different codon *y* is described as,

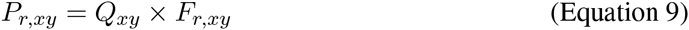

where *Q*_*xy*_ is the rate of mutation from x to y, and *F*_*r,xy*_ is the probability that a mutation from x to y at site r is selected and reaches fixation. In this equation, the mutation rates *Q*_*xy*_ are assumed to be identical across sites whereas the selection is modeled as site-specific and site-independent. The site-specific fixation probabilities *F*_*r,xy*_ were calculated from the experimentally measured amino-acid preferences using the relationship proposed by Halpern and Bruno (Halpern and Bruno, 1998; Bloom, 2014b). The four mutation rate free parameters and the stringency parameter were defined as in (Bloom, 2014b).

We then calculated the phylogenetic likelihood of the observed nucleoprotein sequences given the resulting experimental substitution model *P*_*r,xy*_, the nucleoprotein phylogenetic tree, and the model parameters. The tree consisted of influenza nucleoproteins from either human, swine, equine, or avian hosts. While holding the tree topology fixed, tree branch lengths, and any other model parameters (discussed below), were optimized by maximum likelihood.

To compare overall phylogenetic likelihoods calculated under various substitution models, we calculated the difference in the Akaike Information Criteria (Δ*AIC*) between models. We compared sitespecific models derived from experimentally determined amino-acid preferences to a non-site-specific model. We tested separate site-specific models using the amino-acid preferences from PR/1934 and Aichi/1968. The Aichi/1968 preferences were an average of the amino-acid preferences from the *current study* and *previous study*. In addition, we tested a site-specific model where we combined data from the separate Aichi/1968 and PR/1934 mutational-scanning experiments, by averaging amino-acid preferences for each amino acid at each site across the two homologs, weighting each homolog equally.

The non-site-specific model used the Goldman-Yang (GY94) codon substitution model (Goldman and Yang, 1994), with nucleotide equilibrium frequencies calculated by the CF3x4 method (Pond et al., 2010). In this model, the transition-transversion ratio was optimized by maximum likelihood, along with the mean and shape parameters describing gamma distributions of the nonsynonymous-synonymous ratios (Yang et al., 2000) and the substitution rates (Yang, 1994) across sites. Each gamma distribution was discretized with four categories. In previous comparisons of non-site-specific models, this non-site-specific model performed better than other variants of the GY94 model (Bloom, 2014a,b). All analyses were performed using the software packages `phyloExpCM` (Bloom, 2014a) and `HyPhy` (Pond et al., 2005), and the data, scripts, and descriptions to replicate the results in this article are available at https://github.com/mbdoud/Compare-NP-Preferences.

#### Phylogenetic trees for different influenza hosts

We built phylogenetic trees for nucleoprotein coding sequences from strains of human influenza, swine influenza, equine influenza, and avian influenza. Full-length nucleoprotein sequences were downloaded from the Influenza Virus Resource (Bao et al., 2008), and for each host, a small number of unique sequences per year per influenza subtype were retained. For human influenza, we retained one sequence every other year from each of the H1N1, H2N2, and H3N2 lineages. For swine influenza, we retained one sequence per year from either the North American Classical H1N1 lineage or the Eurasian H1N1 lineage. For equine influenza, we retained one sequence per year from the H3N8 lineage. For avian influenza, one sequence every other year per subtype was retained, and the examined hosts were further restricted to only duck species, to make a sequence set with a size manageable for phylogenetic modeling.

Sequences from each host were aligned by EMBOSS `needle` (Rice et al., 2000), and maximum-likelihood trees were built by `RAxML` (Stamatakis, 2006). Using these trees and the program `Path-O-Gen` (http://tree.bio.ed.ac.uk/software/pathogen/), we identified and removed any sequences that were noticeable outliers from the molecular clock. The final tree contained 37, 46, 29, and 24 sequences from human, swine, equine, and avian hosts respectively.

Maximum-likelihood phylogenetic trees were then built from the nucleoprotein sequence alignment using `codonPhyML` (Gil et al., 2013). The GY94 model (Goldman and Yang, 1994) was run using the CF3x4 nucleotide equilibrium frequencies (Pond et al., 2010) along with maximum-likelihood optimization of a transition-transversion ratio and of a mean and shape parameter describing a gamma distribution of nonsynonymous-synonymous ratios (Yang et al., 2000). This gamma distribution was discretized with four categories. The final, unrooted tree was visualized with `FigTree` (http://tree.bio.ed.ac.uk/software/figtree/) and rooted using the avian clade (Worobey et al., 2014).

## Acknowledgments

We thank Hugh Haddox, Alistair Russell, and Heather Machkovech for critical reading of the manuscript and Trevor Bedford for helpful discussions about statistical analysis. We thank the Summer Institute in Statistics and Modeling in Infectious Diseases at the University of Washington for helpful instruction and the Genomics Shared Resource at the Fred Hutchinson Cancer Research Center for performing high-throughput sequencing. This work was supported by the National Institute of General Medical Sciences of the National Institutes of Health (grant number R01 GM102198). M.D. was supported by NIH Training Grant T32 AI083203 and a fellowship from the Seattle Chapter of the Achievement Rewards for College Scientists Foundation. O.A. was supported by a PhRMA Postdoctoral Fellowship in Informatics.

---

**Supplementary table 1:** Combining experimentally informed substitution models for swine influenza NP. This table differs from Table 1 in that the phylogenetic fit is for the tree of swine NPs shown in Figure 1.

| model | $\Delta\text{AIC}$ | log<br>likelihood | parameters<br>(optimized +<br>empirical) | optimized parameters |
| --- | --- | --- | --- | --- |
| Aichi/1968 + PR/1934 | 0.0 | -6832.4 | 5 (5 + 0) | $R_{A \rightarrow G} = 4.9, R_{A \rightarrow T} = 0.9, R_{C \rightarrow A} = 1.4, R_{C \rightarrow G} = 0.1, \beta = 2.8$ |
| PR/1934 | 390.4 | -7027.6 | 5 (5 + 0) | $R_{A \rightarrow G} = 5.1, R_{A \rightarrow T} = 0.9, R_{C \rightarrow A} = 1.4, R_{C \rightarrow G} = 0.1, \beta = 2.1$ |
| Aichi/1968 | 563.3 | -7114.0 | 5 (5 + 0) | $R_{A \rightarrow G} = 5.0, R_{A \rightarrow T} = 0.8, R_{C \rightarrow A} = 1.4, R_{C \rightarrow G} = 0.1, \beta = 2.4$ |
| GY94, gamma $\omega$, gamma rates | 2153.5 | -7901.2 | 13 (4 + 9) | $\kappa = 5.9, \omega \text{ shape} = 0.3, \text{mean } \omega = 0.0, \text{rate shape} = 2.9$ |

**Supplementary table 2:**
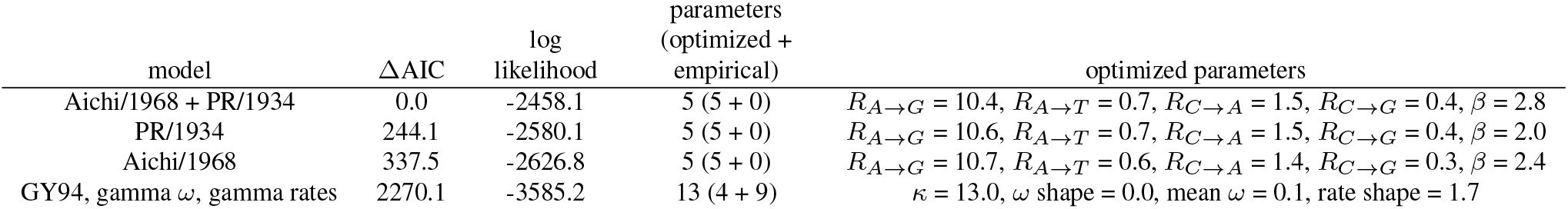
Combining experimentally informed substitution models for equine influenza NP. This table differs from Table 1 in that the phylogenetic fit is for the tree of equine NPs shown in Figure 1.

**Supplementary table 3:**
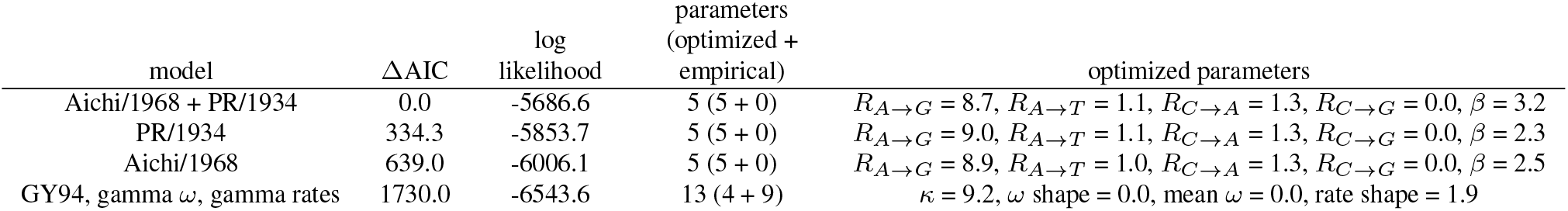
Combining experimentally informed substitution models for avian influenza NP. This table differs from Table 1 in that the phylogenetic fit is for the tree of avian NPs shown in Figure 1.

**Supplementary figure 1:**
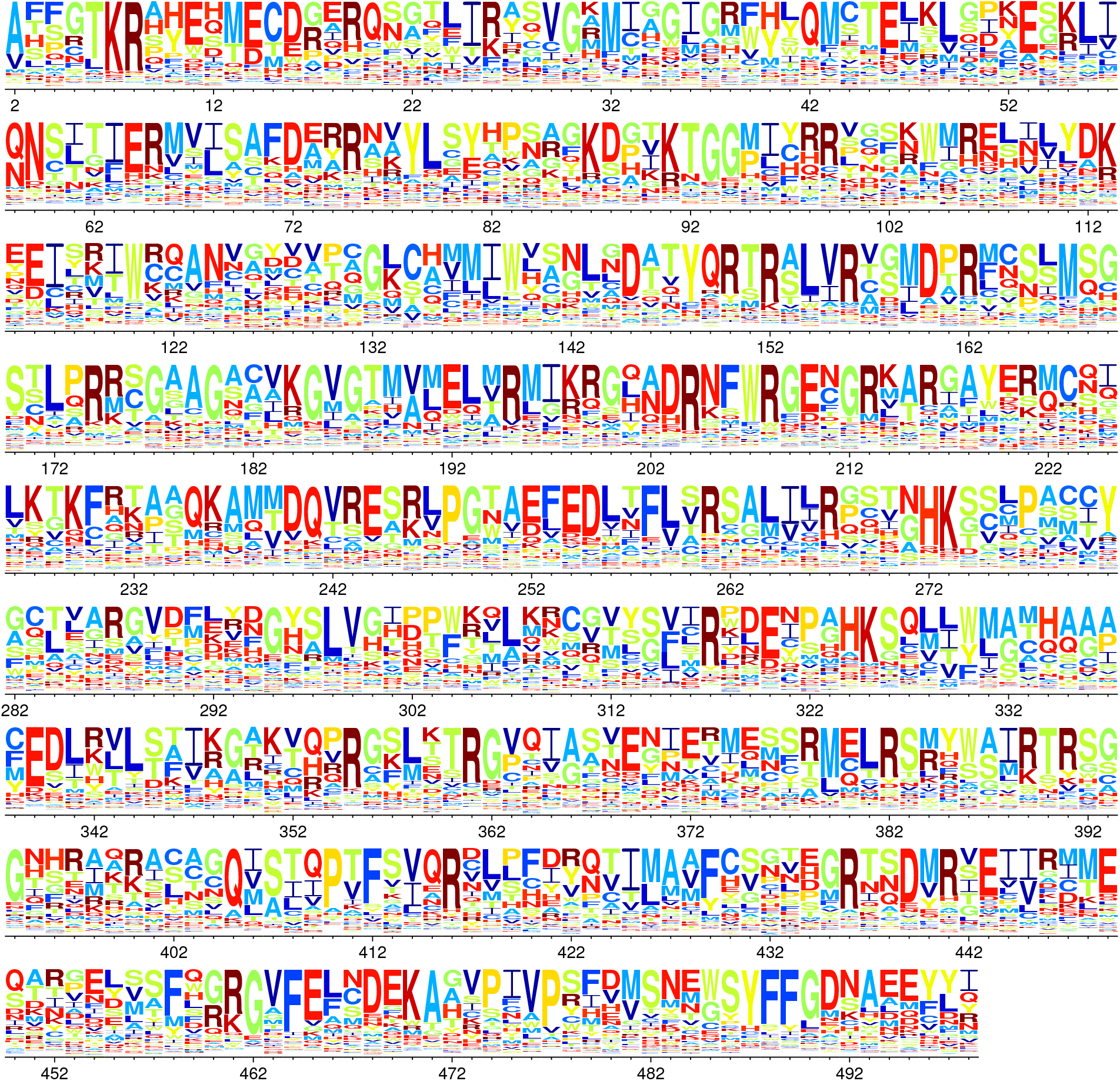
Logoplot of amino-acid preferences for PR/1934 NP. The mean preferences for sites 2 through 498 of PR/1934 are represented in a sequence logo-like visualization created with the program dms_logoplot. The height of each letter is proportional to the preference for that amino-acid at that site.

**Supplementary figure 2:**
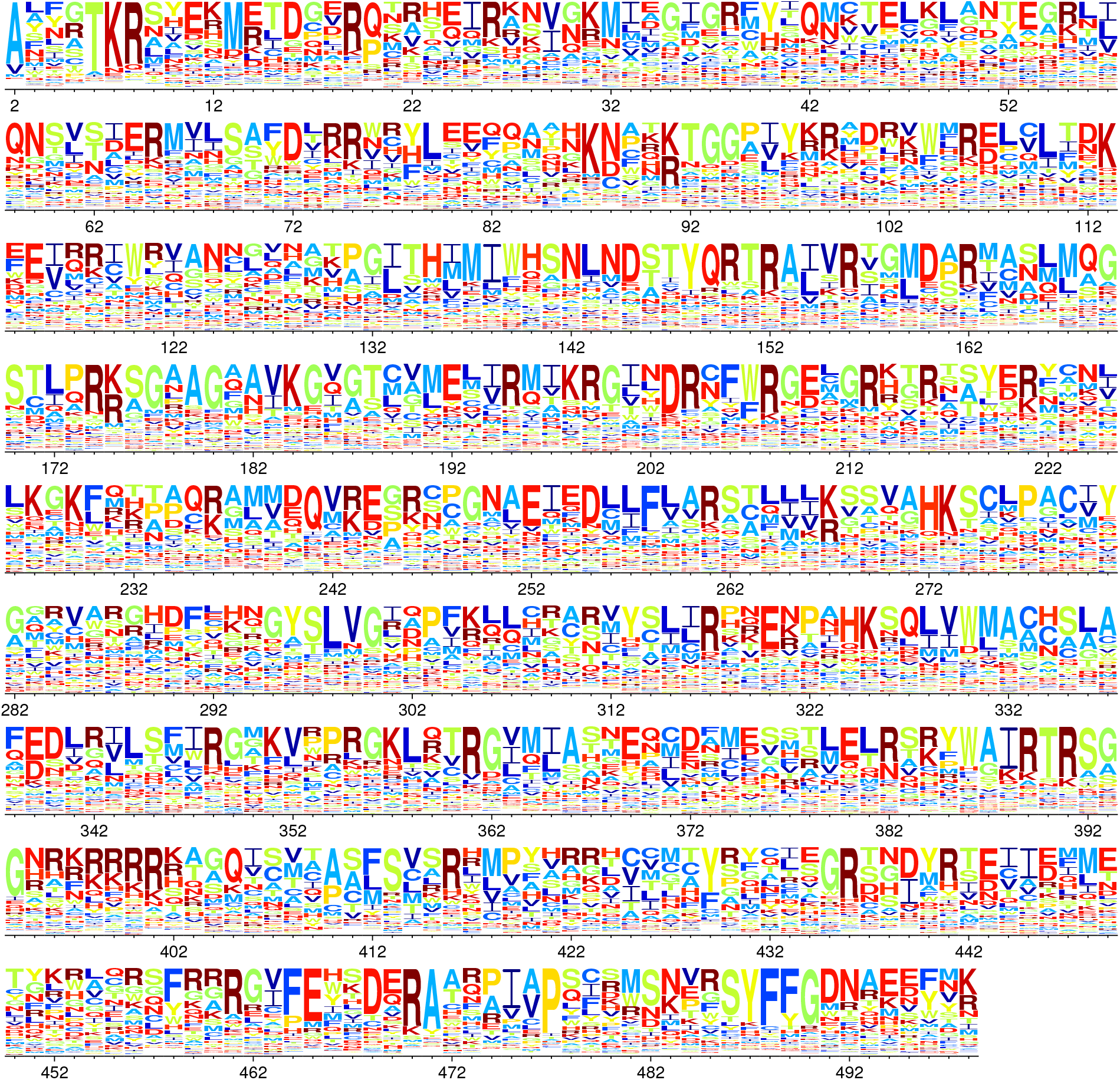
Logoplot of amino-acid preferences for Aichi/1968 NP. The mean preferences for sites 2 through 498 of Aichi/1968 are represented in a sequence logo-like visualization created with the program dms logoplot. The height of each letter is proportional to the preference for that amino-acid at that site.

**Supplementary figure 3:**
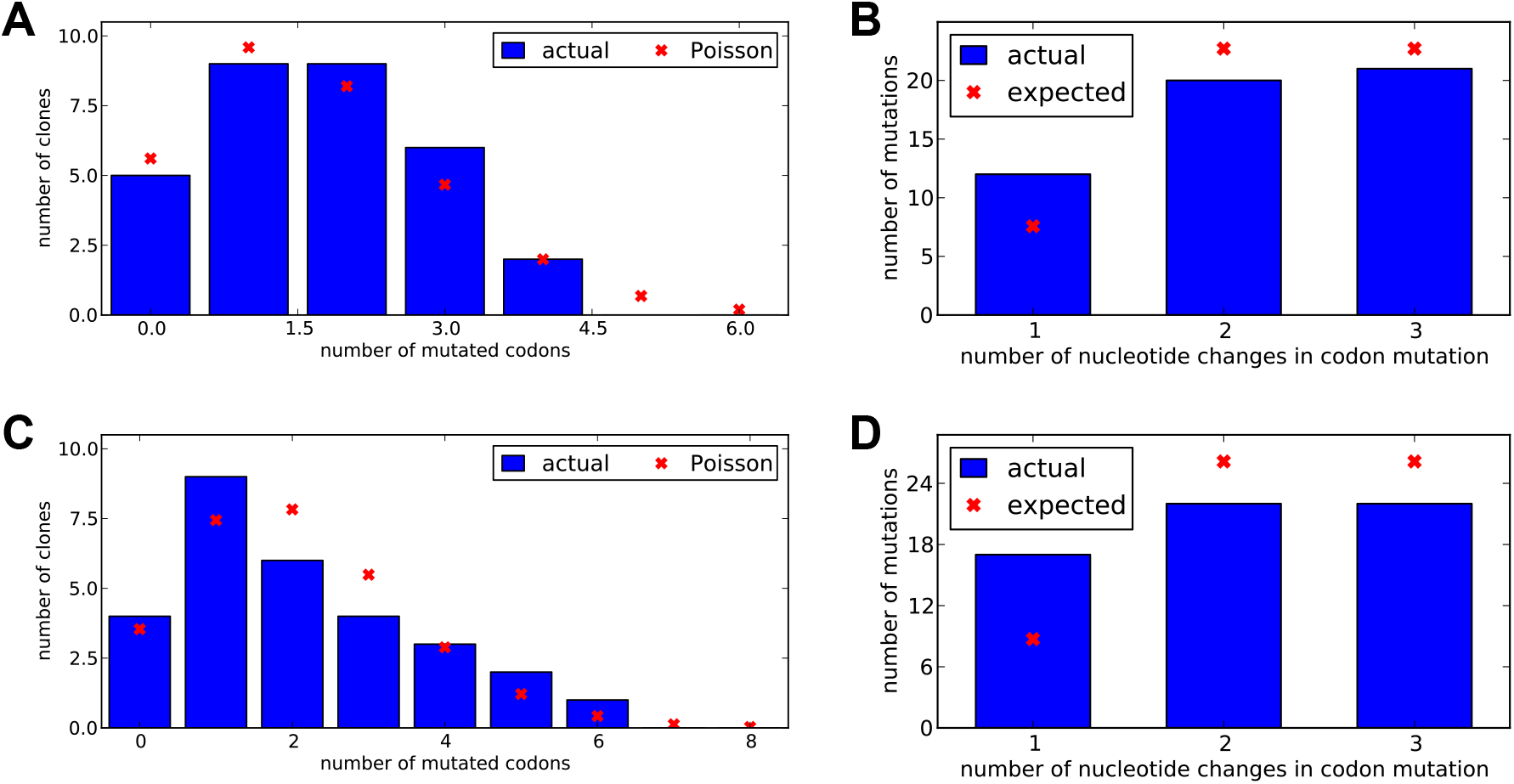
Characterization of plasmid mutant libraries generated by codon mutagenesis. The distributions of number of mutated codons per clone (A, C) and number of nucleotide changes per codon mutation (B, D) were determined by full-length Sanger sequencing of individual clones. A-B: PR/1934 libraries, C-D: Aichi/1968 libraries.

**Supplementary figure 4:**
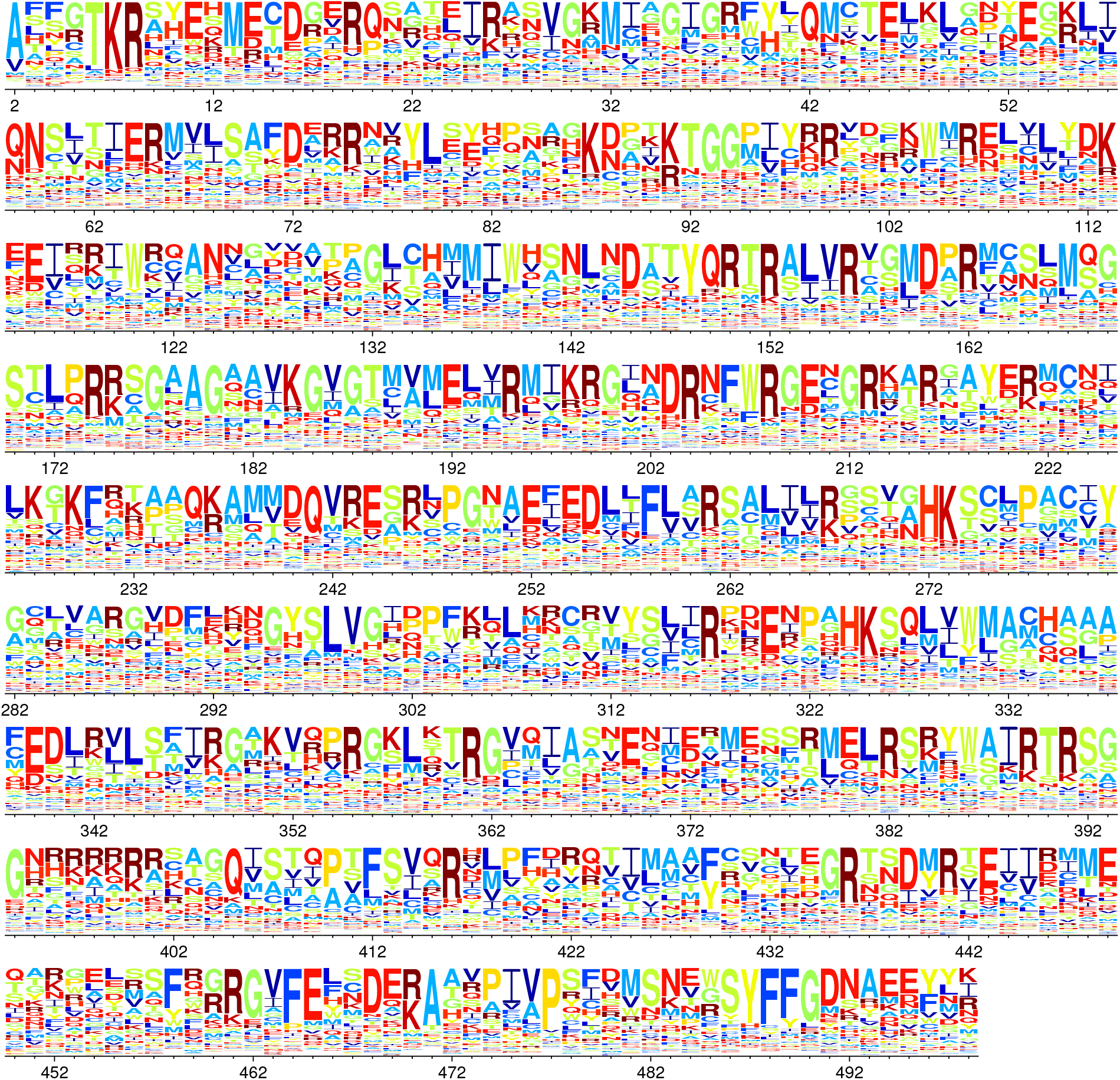
Logoplot of amino-acid preferences for combined PR/1934+Aichi/1968 NP. The mean preferences for sites 2 through 498 of the combined Aichi/1968 + PR/1934 model are represented in a sequence logo-like visualization created with the program dms logoplot. The height of each letter is proportional to the preference for that amino-acid at that site.

**Supplementary figure 5:**
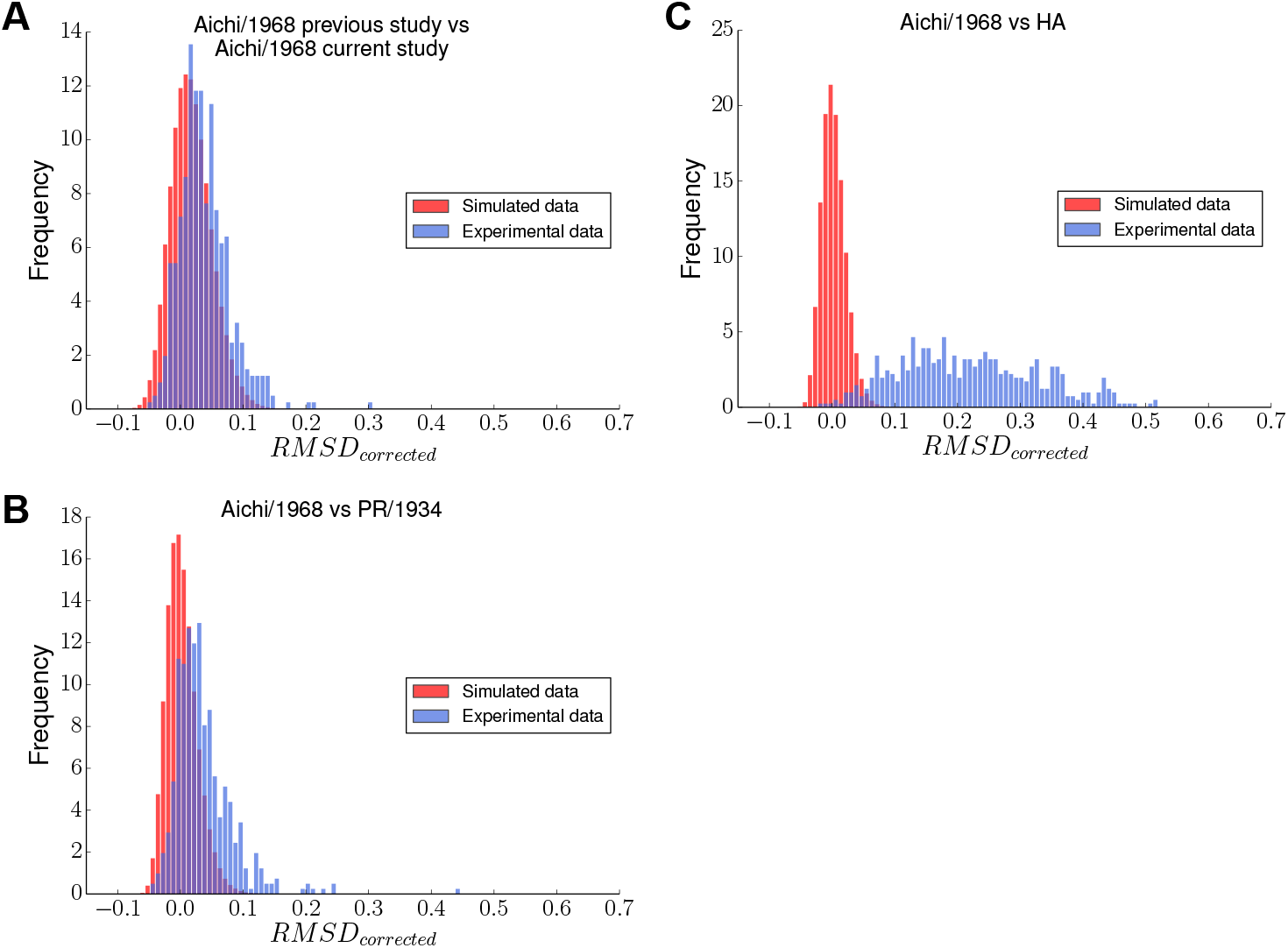
Null distributions of *RMSD_corrected_* generated by simulation. The null distributions generated by simulation are shown in red; experimental distributions are shown in blue.

**Supplementary file 1: Mean amino-acid preferences for PR/1934 NP.** This text file lists the mean amino-acid preferences for sites 2 through 498 in PR/1934 NP. The amino-acid preferences inferred from three biological replicates for PR/1934 NP were averaged at each site.

**Supplementary file 2: Mean amino-acid preferences for Aichi/1968 NP.** This text file lists the mean amino-acid preferences for sites 2 through 498 in Aichi/1968 NP. The average is taken from the average across the previous study replicates and the average from the current study replicates.

**Supplementary file 3: Mean amino-acid preferences for combined PR/1934+Aichi/1968 NP.** This text file lists the amino-acid preferences averaged evenly across the two homologs.

**Supplementary file 4: NP sequence alignment used to build phylogenetic tree.** The alignment consists of human, swine, equine, and avian NP-coding sequences.

**Supplementary file 5: Amino-acid preference RMSD calculations for PR/1934 vs. Aichi/1968 NP.** This table lists summary findings for the comparison of amino-acid preferences between PR/1934 and Aichi/1968 NP. For sites 2 through 498 (the initiating methionine was not mutagenized in our experiments) the amino-acid identity is noted to either be conserved or variable between PR/1934 and Aichi/1968 NP homologs. For variable sites, the PR/1934 amino-acid identity is listed first. The *RMSD_between_*, *RMSD_within_*, and *RMSD_corrected_* statistics calculated for the comparison between 3 replicates of PR/1934 and 10 replicates of Aichi/1968 are listed in the following columns. The next two columns denote sites where the null hypothesis that preferences are identical across homologs can be rejected (under a false discovery rate of 5%) using exact randomization or simulation to generate null distributions of *RMSD_corrected_*, respectively. The final two columns list the mean amino-acid preferences for the most preferred amino acids as measured in each homolog. Amino acids are listed in their order of preference to account for the top 65% of preferences (preferences sum to 1 at each site), with up to three listed for each homolog; full amino-acid preference data is available in Supplementary file 1, Supplementary file 2, and Supplementary file 3.

